# Ketomimetic Nutrients Trigger a Dual Metabolic Defense in Breast Cancer Cells

**DOI:** 10.1101/2024.07.03.601966

**Authors:** Mohini Kamra, Yuan-I Chen, Paula C. Delgado, Erin Seeley, Stephanie K. Seidlits, Hsin-Chin Yeh, Amy Brock, Sapun H. Parekh

## Abstract

While the triggers for the metastatic transformation of breast cancer (BC) cells remain unknown, recent evidence suggests that intrinsic cellular metabolism could be a crucial driver of migratory disposition and chemoresistance. Aiming to decode the molecular mechanisms involved in BC cell metabolic maneuvering, we study how a ketomimetic (ketone body-rich, low glucose) medium affects Doxorubicin (DOX) susceptibility and invasive disposition of BC cells. We quantified glycocalyx sialylation and found an inverse correlation with DOX-induced cytotoxicity and DOX internalization. These measurements were coupled with single-cell metabolic imaging, bulk migration studies, along with transcriptomic and metabolomic analyses. Our findings revealed that a ketomimetic medium enhances chemoresistance and invasive disposition of BC cells via two main oncogenic pathways: hypersialylation and lipid synthesis. We propose that the crosstalk between these pathways, juxtaposed at the synthesis of the glycan precursor UDP-GlcNAc, furthers advancement of a metastatic phenotype in BC cells under ketomimetic conditions.

## INTRODUCTION

Breast cancer (BC) represents the leading cause of cancer-related deaths among women worldwide.^1^ It is widely recognized for its strong propensity to metastasize to bone, liver, and the brain. In the process of metastasis, mammary epithelial cells (MECs) undergo a transformation, adopting an invasive phenotype, and migrate away from the primary tumor to infiltrate and establish themselves elsewhere in the body.^2,3^ Unfortunately, the triggers for the transformation of MECs remain unknown and recent evidence has suggested that switches in metabolic activity are initiators for invasive and drug-resistant phenotypes.^4^

Metabolic manipulation is a well-known hallmark of cancer. Typical cellular metabolism uses oxidative phosphorylation (OXPHOS) for ATP generation. However, tumor cells are also supported by an additional aerobic glycolytic pathway to meet their increased energetic demands, a process known as the Warburg effect, i.e. lactate formation and shuttling NADH into the mitochondria for ATP generation.^5^ This facet of cancer cells, along with additional alterations in lipid metabolism, together regulate cancer aggressiveness and therapy resistance.^6,7^ Particularly, tumors in the breast are surrounded by bulk adipose tissue that provides a steady supply of lipids, which are the metabolic precursors for beta-oxidation, a supporting pathway for extra energetic demands of cancer cells.^8,9^ It is believed that the antioxidant and the buffering environment created by lipid synthesis/metabolism protects the cells against the chemotherapeutic-induced ferroptosis.^10,11^ Another crucial pathway is the mevalonate (MVA) biosynthetic pathway, which is associated with reprogramming of lipid metabolism in specific cancer cells.^12^ The MVA pathway is essential for ensuring supply of cholesterol-derived metabolites for tumor propagation and is also believed to be linked to chemoresistance.^13^ Moreover, mammary adipocytes have been shown to stimulate BC invasion through fatty acid-mediated metabolic remodeling.^14,15^ Increased reliance of BC cells on fatty acids from surrounding adipose tissue has also been seen to promote cancer invasiveness through the epithelial-to-mesenchymal transition (EMT).^16,17^ During EMT, BC cells acquire an increased mesenchymal phenotype and show enhanced migratory capacity and elevated rates of metastasis.^18^

One more interesting property of BC cells is hypersialylation, which is the presence of increased sialic acid residues on membrane-associated glycoproteins and glycolipids, has been established as an oncogenic biomarker by numerous studies over the recent decades.^19–21^ Studies have highlighted the vital role of glycocalyx sialylation in several physiological processes,^22^ including their participation in cell-cell and cell-matrix interactions.^23^ In BC, pronounced expression of sialoglycans correlates with the aggressiveness of the tumor and its capacity to invade neighboring tissue.^24^ Recent research has focused on the development of strategies to inhibit aberrant sialylation as a potential route to control tumor growth and promote immune response while also enhancing the effectiveness of conventional chemotherapeutics and contemporary immunotherapeutics.^25–28^ As hypersialylation relies heavily on enhanced sialic acid biosynthesis from the oncogenic glycolytic upsurge in cancer cells, it stands to reason that nutrient (dietary) changes could impact the extent of sialylation of the glycocalyx. The link between diet and sialylation has been studied in the context of aging, and high fat diet or dietary restrictions have been shown to alter the acetylation pattern of cell surface sialic acids, based on the availability of acetyl CoA.^29^ It has also been observed that exogenously fed sialic acid attenuates inflammation and oxidative stress induced by a high-fat diet.^30^ Independently, very recent studies have begun examining the effect of different diets on the efficacy of anti-cancer therapy in animal models;^31^ however, no studies have probed the direct effects of ketomimetic nutrient medium on glycocalyx sialylation that may mediate changes in therapy success or invasive disposition of BC cells.

Although recent research has shown that increased dietary lipids can act as a chemotherapy adjuvant in many cancer models,^32,33^ there are mixed findings in animals adopting a ketogenic diet with both enhancement and reduction of chemoresponse observed.^31,34,35^ This discrepancy may be dependent on the specific tumor model being used. More importantly, these studies are mostly based on a macroscopic examination of animal survival or tumor size over time. A connection between local metabolic fuel perturbation, cellular/tumor metabolism, and chemotherapy was recently reported,^36^ but unfortunately the connections to specific mechanisms are unclear. Our report is the first one to offer a clear mechanistic understanding of the effect of local metabolic fuel on the key drivers of BC metastasis and chemoresistance. Herein, we investigate the crosstalk of these two key oncogenic drivers, lipid accumulation and hypersialylation, and their collective response to ketone bodies as alternate nutrient sources available to BC cells. We investigate the direct role of a ketomimetic nutrient medium (rich in ketone bodies and low in glucose) on BC cell aggressiveness, invasiveness, and therapy resistance coupled with metabolic signatures.

## RESULTS AND DISCUSSION

### Ketomimetic nutrient medium enhances glycocalyx sialylation and confers chemoprotection to BC cells

Sialic acids, being correlated with increased malignancy, make the hypersialylation of cancer cells a potential avenue by which to assess tumorigenicity and, possibly, invasive disposition.^21^ We used the bioorthogonal click chemistry imaging technique to measure degree of glycocalyx sialylation with the azido mannose, Ac_4_MaNAz, using the copper-free methodology developed by Bertozzi et al.^37^ To quantify relative sialylation at the membrane, we counter-stained the plasma membrane and calculated the integrated fluorescence intensity/ perimeter of membrane. This further ensures that larger and smaller cells are quantified without bias. Our data revealed that the invasive MDA-MB-231 BC cell line boosted its sialic acid levels by ∼ 2-fold when glucose in their nutrient medium is lowered to 1 g/L from 4.5 g/L and when ketone body forming 3-hydroxybutyrate (HB) supplemented low glucose medium is used (**Figure 1 A and B**).

**Figure 1.**
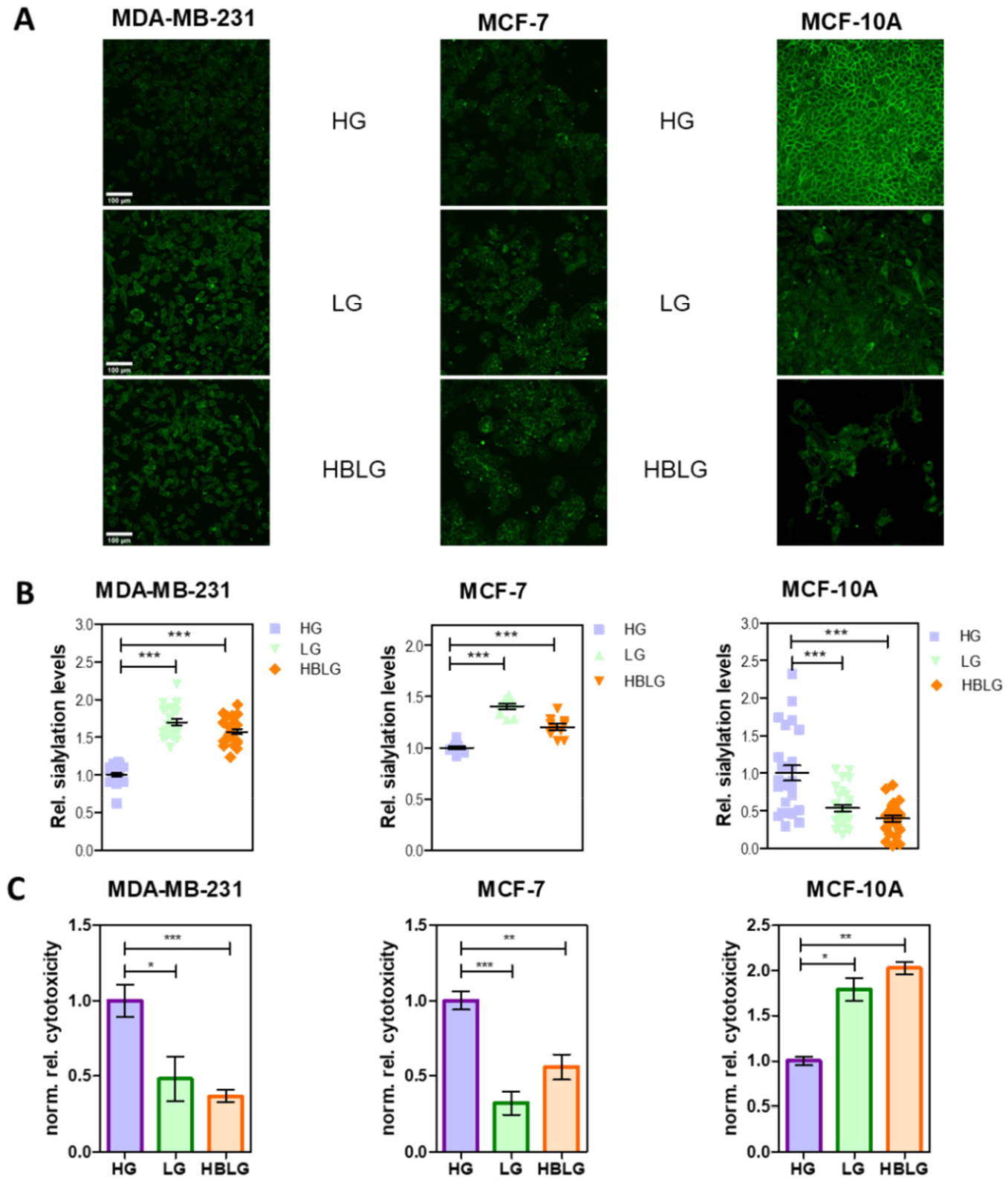
BC cell sialylation and susceptibility to chemotherapy is nutrient dependent and internally inversely related. (A) Representative confocal microscopy images of MDA-MB-231, MCF-7 and MCF-10A cells obtained using azide-alkyne click-chemistry following metabolic labeling of membrane sialic acids in the respective nutrient media: high glucose (HG), low glucose (LG) and HBLG (3-hydroxybutyrate-containing, low glucose). Green: AF488; scale bar: 100 µm. All images were collected at identical conditions and are displayed with the same LUT. (B) Relative sialylation levels in each cell line normalized to HG control as quantified using AF 488 intensity/perimeter of membrane in an image of labeled cells. For each cell type, images from two separate experiments (N=2) were used with 10-15 different FOVs collected for each experiment. A custom pipeline created in Cell profiler was used for quantification based on generation of membrane mask using an image of the cells with CellBrite Membrane stain. Scatter plots shown here with black horizontal lines representing means, and error bars representing the standard error of mean (SEM), *p*-values were calculated using a two-tailed unpaired t-test in GraphPad Prism. (C) Normalized relative cytotoxicity of DOX (1µM) in MDA-MB-231, MCF-7 and MCF-10A cells in the respective nutrient media plotted using mean values were error bars represent SEM and *p*-values were computed using a two-tailed unpaired t-test in GraphPad Prism. *, **, *** indicate p<0.05, p<0.01, and p<0.001, respectively.

Switching the medium to low glucose (LG) or HB-rich low glucose (HBLG) media allowed us to explore how changing nutrients affect sialylation. The effect of supplementation of HB on sialylation when compared to LG alone was not significant, and both conditions showed more sialylation than the standard high glucose (HG) medium. In the MCF-7 (somewhat less invasive) BC cell line, switching to LG alone caused a 1.5-fold increase in sialylation compared to HG, and HBLG marginally attenuated the effect of low glucose. Meanwhile, non-cancerous MECs, the MCF-10A cell line showed a significant reduction in sialylation as the amount of glucose was lowered and exhibited a further drop to less than 0.5-fold in the HBLG medium compared to the HG medium. The opposite impact of nutrient switching on sialylation of cancer versus non-cancer cells is intriguing. From these results, we infer that: (a) cell nutrient sources can manipulate cellular metabolic pathways to modify their protective glycocalyx and (b) the differential metabolic characteristics of cancer vs. non-cancer cells have direct consequences on their ability to sialylate their respective surfaces. Although hypersialylation is considered an indication of enhanced chemoprotection and metastatic potential,^38^ the molecular mechanisms connecting elaboration of sialoglycans and efficacy of chemotherapeutic response remain unclear. As can be seen from **Figure 1C**, LG and HBLG media progressively decrease the sensitivity of the BC cells (MDA-MB-231 and MCF-7) to the chemotherapeutic agent Doxorubicin (DOX).

The presence of ketone body-forming metabolites in the media appeared to protect the BC cells from the chemotherapeutic agent. In contrast, both LG and HBLG media sensitized the non-cancerous MEC’s, MCF-10A cells to the cytotoxic action of DOX, as confirmed by a lactate dehydrogenase (LDH) cytotoxicity assay. In addition to the opposing effects of nutrient medium in case of non-cancerous and BC cells, we also observed an inverse correlation between the trends of sialylation and DOX-mediated cytotoxicity. That is, enhancement in glycocalyx sialylation suppressed DOX-induced cytotoxicity and *vice versa*.

### Ketomimetic nutrient medium promotes a glycolytic phenotype

Label-free metabolic imaging using fluorescence lifetime microscopy (FLIM) of NADH autofluorescence was used to assess changes in cellular metabolic phenotype, in real-time, in response to drug treatments in the different nutrient media. **Figure 2A** shows representative images of the NADH intensity and NADH metabolic index in MDA-MB-231 and MCF-10A cells. In general, HBLG medium promotes glycolysis in both the cell types, non-cancerous and invasive BC (**Figure 2B, C**), referring to MCF-10A and MDA-MB-231 cell lines, respectively. Interestingly, DOX treatment causes a progressive enhancement in the glycolytic preference (**Figure 2C**) over time. Notably, while a comparable increase in glycolytic activity is noted in MCF-10A and MCF-7 cells in HBLG media following DOX treatment, MDA-MB-231 cells do not show enhanced glycolysis with DOX treatment. This suggests that either the suppression of the DOX response or the inherent glycolytic activity of MDA-MB-231 cells is maxed out in LG and HBLG media. Images of NADH intensity and metabolic index for all the three FOV’s in each condition are provided in **Supplementary Figures S1-S6**.

**Figure 2.**
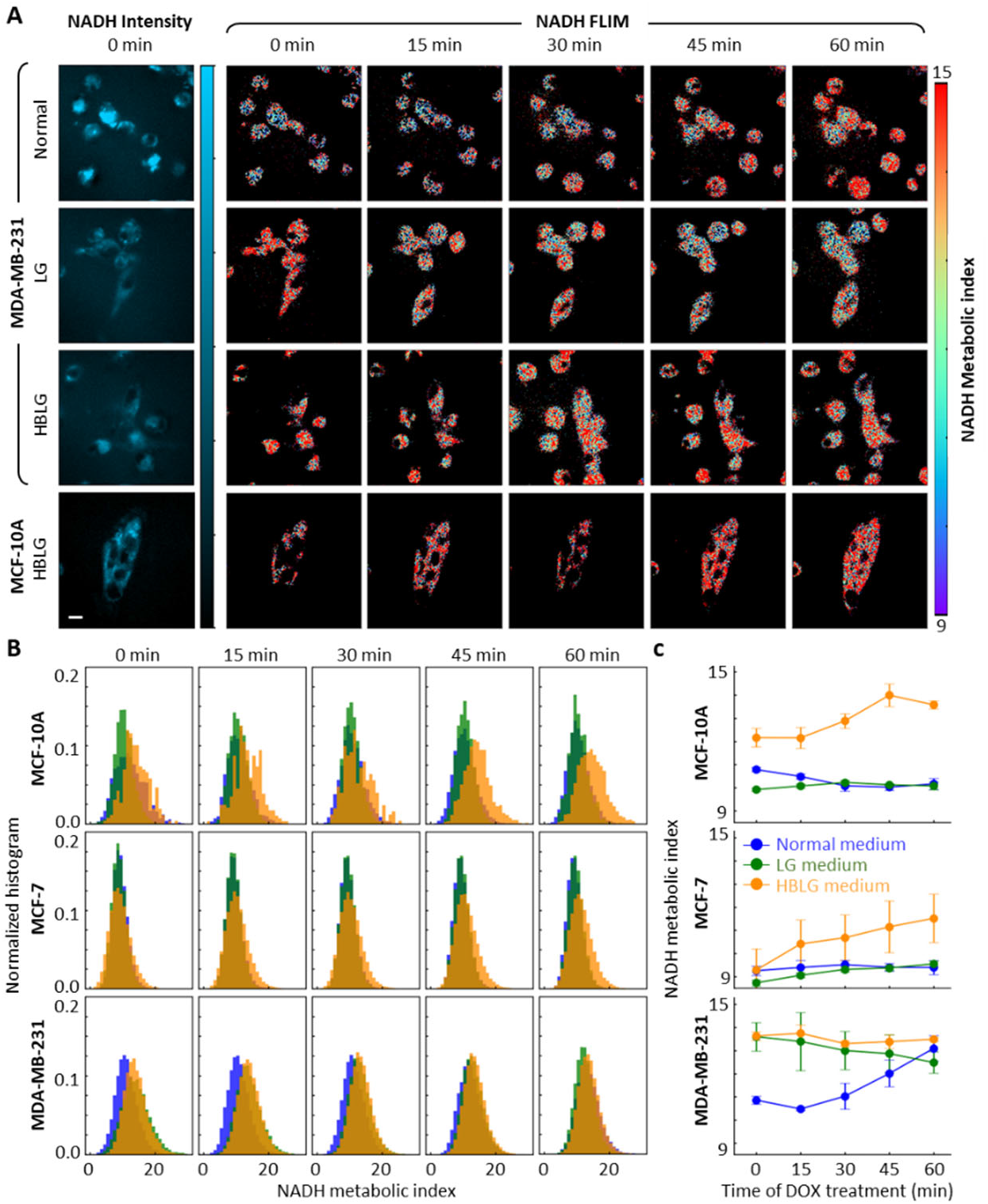
Alterations in NADH metabolic index of cells cultured in different media during the first hour of DOX treatment. (A) Upper three panels: representative intensity contrast images of MDA-MB-231 cells under normal, LG, and HBLG medium at 0 min are shown on the left and FLIM images (α1/α2) are shown towards the right in the order 0 – 60 min. Scale bar, 10 µm. Similarly representative images of MCF-10A cells in HBLG are shown in the lower panel. (B) Normalized NADH metabolic index histograms obtained from MCF-10A, MCF-7, and MDA-MB-231 in normal, LG, and HBLG medium over time. (C) Averaged NADH metabolic index obtained from MCF-10A, MCF-7, and MDA-MB-231 in normal, LG, and HBLG medium over time. Circular symbols are median values and error bars represent standard deviations over all pixels from a set of three different FOV’s per condition.

A general belief is that preference for glycolysis over oxidative phosphorylation (OXPHOS) in cancer cells is indicative of a stronger propensity for invasion rather than proliferation. This hypothesis was further confirmed using functional assays of proliferation and migration (discussed below). Whether the increase in glycolytic phenotype in a non-cancerous MEC line, like MCF-10A, translates into a slower or a faster proliferation rate, is an interesting question that we explore below.

When glucose levels were altered in the media, total cellular glucose consumption for BC cells remained constant (**Figure 3A**). Moreover, the addition of HB to LG medium did not change glucose uptake appreciably. Thus, we can conclude that reducing glucose availability does not affect total glucose uptake. We observed that, as opposed to non-cancerous breast epithelial cells, glucose deprivation makes both the BC cell lines: MDA-MB-231 and MCF-7 more “glucose-hungry”. These BC cells consume nearly 3-4 times higher percentage of available glucose (**Supplementary Figure S7**) to maintain the same total amount of uptake. Interestingly, the same total uptake is maintained despite the lack of insulin in LG and HBLG media. This is consistent with the enhanced glycolytic phenotype in low glucose nutrient media observed from FLIM (NADH metabolic index at time 0).

**Figure 3.**
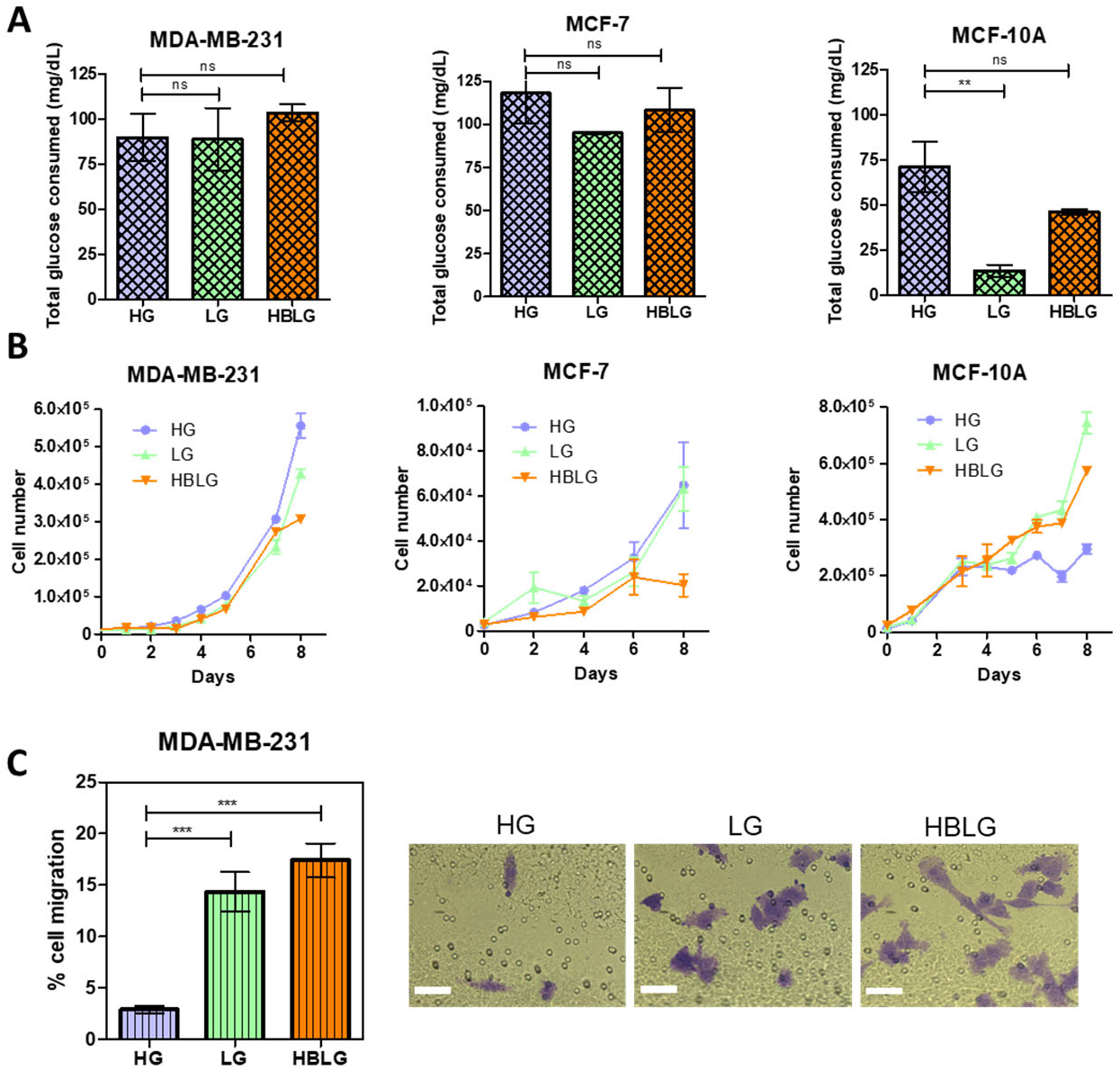
Glucose consumption and migratory propensity of cells in different nutrient media. (A) Glucose consumption patterns of MDA-MB-231, MCF-7 and MCF-10A cells in the respective nutrient media: high glucose (HG), low glucose (LG) and HBLG (hydroxybutyrate containing low glucose). Changes in the amount of glucose consumed by the cells after four days of culture are plotted with data from n=3 and N=2 being pooled together. GraphPad Prism was used for plotting the data with colored bars representing means and error bars representing SEM, two-tailed unpaired t-tests were performed for column-wise statistical analysis. (B) Cellular proliferation profiles of the three cell lines in the respective nutrient media. Symbols denote mean values and error bars denote SEM from N = 2 and n = 2. (C) Cellular migration profile of MDA-MB-231 cells in response to changes in nutrient media quantified using images of crystal violet stained cells that have migrated across transwell membrane. Images from two separate experiments (N=2) were used with 8-10 different FOVs collected for each experiment. The percentage (%) occupied area was measured using ImageJ software; bar graph displays mean values with error bars denoting SEM. Representative images are shown in the right panel, scale bar = 50 µm. ns, **, *** indicate p>0.05, p<0.01, and p<0.001, respectively.

### Ketomimetic nutrient medium slows proliferation and promotes migration of BC cells

As measured from an Erythromycin B-based cell counting assay, it was observed that the ketomimetic nutrient medium lowered the proliferation rates of MDA-MB-231 and MCF-7 BC cells (**Figure 3B**). Interestingly, glucose deprivation and ketone body supplementation led to an increase in the proliferation rate of non-cancerous MCF-10A cells, which may explain their chemosensitization (since DOX primarily acts during cell division). Thus, we suggest that the enhanced glycolytic phenotype in MCF-10A cells in response to ketomimetic medium translated into enhanced proliferative abilities, rendering them more susceptible to the antineoplastic agent DOX. On the other hand, BC cells that showed decreased proliferation in ketomimetic medium might be employing it as a mechanism to escape the antineoplastic effects of DOX, rendering decreased cytotoxicity. If indeed the BC cells fed with ketone body-rich media use reduced proliferation as a mechanism of chemoprotection, despite consuming the same amount of glucose, this raises the following question: what is the consumed glucose being used for, if not for proliferation in the BC cells? This is especially interesting as the BC cells are consuming the same amount of glucose even after reducing the available amounts of glucose in the media and after ketone body supplementation in the LG media. The answer to this question lies, at least partially, in the sialylation data. It appears that one main sink where the incoming glycolytic flux was channeled, was the sialic acid biosynthesis pathway.^31^ Our observations of sialylation (**Figure 1**), glycolysis (**Figure 2, 3A**), and proliferation (**Figure 3B**) are consistent with this hypothesis and suggest that increased glycolysis feeds hypersialylation in a ketomimetic medium, which can be rationalized via the hexosamine biosynthetic pathway (HBP). HBP is responsible for shunting of a majority of fructose-6-phosphate (F-6-P) formed from glucose to the N- and O-glycosylation pathways, including sialic acid synthesis and membrane decoration.^39^

Since the proliferative abilities of the BC cells are inversely affected by the presence of ketone bodies, we wondered if BC cells showed differences in migration. A difference in migration in BC cells with ketogenic media is further suggested since sialylation is increased, which has also been linked to metastasis.^24^ A transwell migration assay demonstrated that MDA-MB-231 subjected to LG and HBLG media had increased migratory ability compared to those cells in the standard, HG media (**Figure 3C**). The increased migratory ability of these cells suggests enhanced metastatic potential. Due to inherently poor migratory tendencies of MCF-7 and MCF-10A cell lines,^40,41^ these cells showed undetectable chemotactic migration across the membrane of the transwell inserts.

### Transcriptomic analysis confirms the mechanism of differential responses of cancer vs. non-cancer cells

The opposite response of non-cancerous and BC cells to the same alteration in nutrient medium could be attributed to the lipid-processing machinery of cancer cells. Normal cells have a tightly regulated machinery for synthesis of the fatty acid and cholesterol, which becomes dysregulated only upon oncogenic transformation.^12^ Another metabolic switch that happens as a result of nutrient sensing by cancer cells is activation of the HBP, which involves the conversion of F-6-P to UDP-GlcNAc and further sialylation.^42^ We used rt-*q*PCR to confirm that the mechanistic drivers of these pathways were differentially expressed in in malignant and non-malignant MECs. We probed the expression levels of select crucial enzymes involved in HBP and sialylation. Comparison of the fold changes relative to the control, MCF-10A cells, are shown in **Supplementary Figure S8**. Our results show that indeed the expression of *GFPT1* (also called GFAT), the rate-limiting enzyme of HBP, is several folds higher in BC cells, MCF-7 and MDA-MB-231, compared to that in MCF-10A cells. Similarly, the expression of *GNE*, the enzyme that feeds UDP-GlcNAc to the sialic acid biosynthesis is significantly higher in the BC cells than the non-cancerous, MCF-10A cells. Importantly, the prominent sialyl transferase *ST6GAL1* showed a significantly higher expression in BC cells as compared to normal cells. The BC cells also display a higher *FASN* expression level than the non-cancerous cells indicative of a higher propensity to synthesize fatty acids. Thus, it appears that hypersialylation and lipid synthesis together contribute to the response of BC cells to ketone-body rich nutrient medium.

### Increased sialylation at the glycocalyx directly blocks DOX internalization

Apart from slowed proliferation and enhanced migration, what are the additional ways by which membrane sialylation could contribute to chemoresistance? We hypothesized that additional sialylation of the glycocalyx in BC cells in response to ketomimetic nutrients would trap or sequester DOX to inhibit uptake. Sialic acids are negatively charged, and DOX is positively charged, so such an effect is plausible. It is possible that the hypersialylated and extensively branched glycocalyx brush can electrostatically trap DOX and inhibit its cellular internalization. We measured cellular internalization using fluorescence microscopy of DOX uptake before and after sialidase treatment (to remove sialic acids from the glycocalyx). We first confirmed that sialidase treatment removed sialic acids as expected (**Supplementary Figure S9**). Next, we quantified the fold-change in DOX fluorescence per cell for each of the nutrient conditions in MCF-7 and MDA-MB-231 cells. As can be seen in **Figure 4A, B**, removal of sialic acids from the glycocalyx using sialidase treatment significantly improved DOX uptake in the cells. This observation confirmed our hypothesis and shows for the first time how hypersialylation of the glycocalyx directly obstructs the cellular internalization of DOX. Data for an experiment performed where sialic acid was labeled in the same cells as Dox (for confirmation of sialidase action) is shown in the supplementary information (**Supplementary Figure S10**).

**Figure 4.**
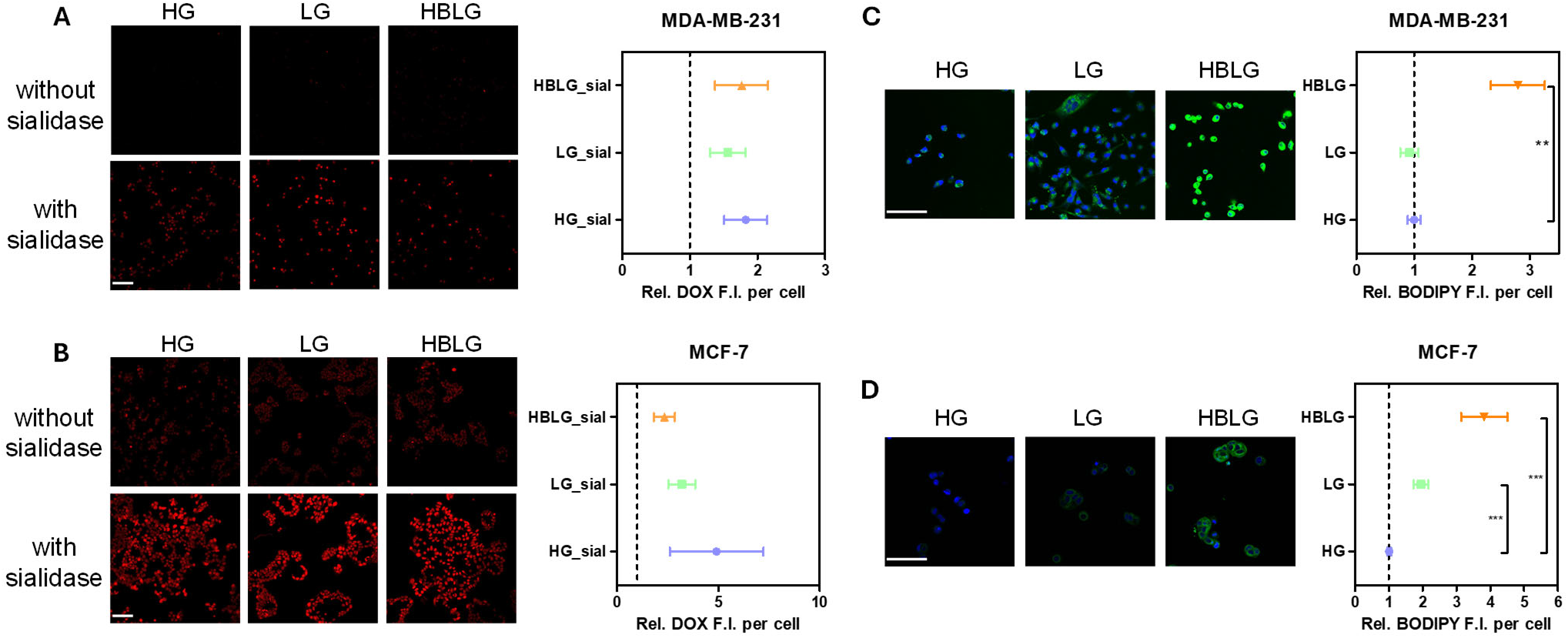
Sialylation and lipid accumulation in ketomimetic media contribute to DOX chemoprotection. (A) Confocal fluorescence imaging of MDA-MB-231 cells obtained after DOX internalization with and without sialidase treatment in the respective nutrient media: high glucose (HG), low glucose (LG) and HBLG (hydroxybutyrate containing low glucose). Red: DOX; scale bar: 100 µm. (B) Confocal fluorescence imaging of MCF-7 cells obtained after DOX internalization with and without sialidase treatment in the respective nutrient media. Red: DOX; scale bar: 100 µm. (C,D) Representative confocal microscopy images of MDA-MB-231 (C) and MCF-7 (D) cells obtained using BODIPY staining for detection of lipid droplets in the respective nutrient media. DRAQ5 was used as a nuclear counterstain. Green: BODIPY 505, Blue: DRAQ5; scale bar: 100 µm. Fluorescence quantification is shown on the right side of each panel. All images were collected at identical conditions and are displayed with the same LUT. The DOX or BODIPY signal per cell was quantified as I_DOX_-I_blank_/cell # or I_Bodipy_-I_blank_/cell #, respectively and fold changes were computed relative to no sialidase and HG as controls, respectively. For each cell type, images from at least three separate experiments (N=3) were used with 10-15 different FOVs collected for each experiment. For all graphs, symbols represent means and error bars denote SEM. **, *** indicate p<0.01, and p<0.001, respectively.

### Nutrient alteration to ketone bodies drastically shifts metabolomic profiles of BC cells

Due to the involvement of lipid metabolism in chemoresistance, we next explored the effect of ketomimetic medium on lipid synthesis and accumulation in the BC cells. As can be seen in **Figure 4C** and **D**, BODIPY 505 staining showed a drastic increase in lipid droplet (LD) accumulation in the BC cells when the nutrient medium was shifted from HG to HBLG compositions. Since the LG medium did not induce LD formation or accumulation compared the HG media, we surmise that ketone bodies in HBLG media feed the MVA and fatty acid synthesis pathways. The non-cancerous cell line, MCF-10A, showed minimal accumulation of LDs in response to ketone body uptake (**Supplementary Figure S11**), suggesting the participation of specialized oncogenic pathways in the processing and storage of ketone bodies in BC cells. Thus, we propose that in a ketone body-rich medium, the cells employ dual mechanisms for drug resistance: sialylation and LD accumulation.

Since the most significant changes in sialylation, migration, and therapeutic resistance were observed in MDA-MB-231 cells, we further investigated their molecular metabolic fingerprint using MALDI mass spectrometry imaging (MALDI-MSI). MALDI-MSI of MDA-MB-231 cells cultured in HG, LG or HBLG media was performed to find metabolomic information. An unsupervised separation of the data using principal component analysis (PCA) is shown in **Figure 5A**. We find that the HG population displays a clearly distinct metabolic profile compared to the LG and HBLG groups. To perform an enrichment analysis, we arranged the metabolites in increasing order of loading on principal component 1 and found that the HG population was enriched in sugar-based metabolites, e.g., sorbitol, mannitol, cellobiose and others (**Figure 5B**).

**Figure 5.**
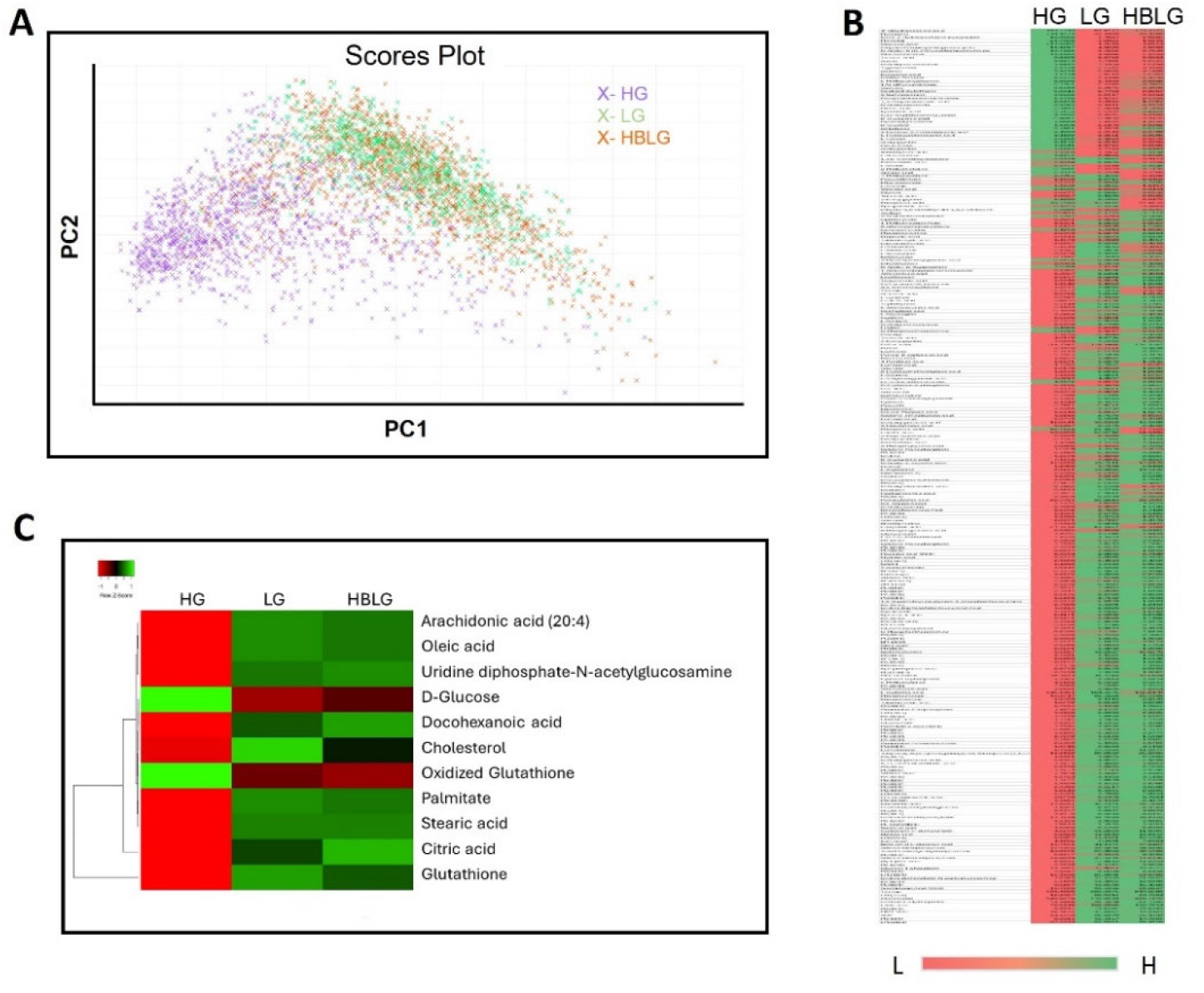
MALDI-MSI based metabolomic profiling of MDA-MB-231 cells in three different nutrient media: HG, LG and HBLG. (A) Principal Component Analysis of MALDI-MSI metabolomic data. (B) Metabolites were arranged in increasing order of loading in PC1, to obtain a metabolite-enrichment based separation. On the color scale bar, L and H denote lowest and highest values respectively, of a particular metabolite across the three nutrient media. (C) Relative abundance of certain key metabolites displayed as a dendrogram-based heatmap.

The LG and HBLG populations, on the other hand, were enriched in UDP-GlcNAc and lipids (cholesterol, stearic acid, oleic acid, and many long chain fatty acids) - fully consistent with our other results (**Figure 4C, D**). A dendrogram-based heatmap of all identified metabolites in negative-ion mode MS is shown in **Supplementary Figure S12** and the relative abundance of key metabolites of interest are displayed as a dendrogram based heatmap in **Figure 5C**. The relative distributions of each of these metabolites with statistical analysis are shown in **Supplementary Figures S13-S24**.

Overall, the MALDI-MSI based metabolomic analysis is reflective of the vast alteration in metabolic signature of the BC cells upon nutrient alteration, which in turn results in phenotypic alterations that lead to enhanced chemoresponse. It is known that UDP-GlcNAc, the product of HBP, is the precursor of most cancer-promoting glycans, including the hypersialylated ones. While a chunk of sialylation occurs via heightened shunting of F-6-P from the elevated glycolysis, ketone bodies can also directly (and independently) form UDP-GlcNAc by generating additional Acetyl CoA. Hence, we can conclude that it is this cross-over of lipid synthesis and glycan synthesis pathways that might be leading to elevation of invasiveness in BC cells in response to ketomimetic media (**Figure 6**). We postulate that the elevated glycolysis in cells grown in ketomimetic media feeds (a) the sialylation pathway via deregulated HBP and (b) the fatty acid synthesis and MVA/cholesterol synthesis pathways, leading to fat storage in LD’s. Functionally, these pathways synergistically promote (i) migratory propensity via sialic acid driven extracellular cues, and (ii) DOX resistance by not only lowering the amount of drug entering the cells but also by promoting de-novo lipogenesis, which is known to confer chemoprotective effect^10,11^ from the drug that has entered the cells.

**Figure 6.**
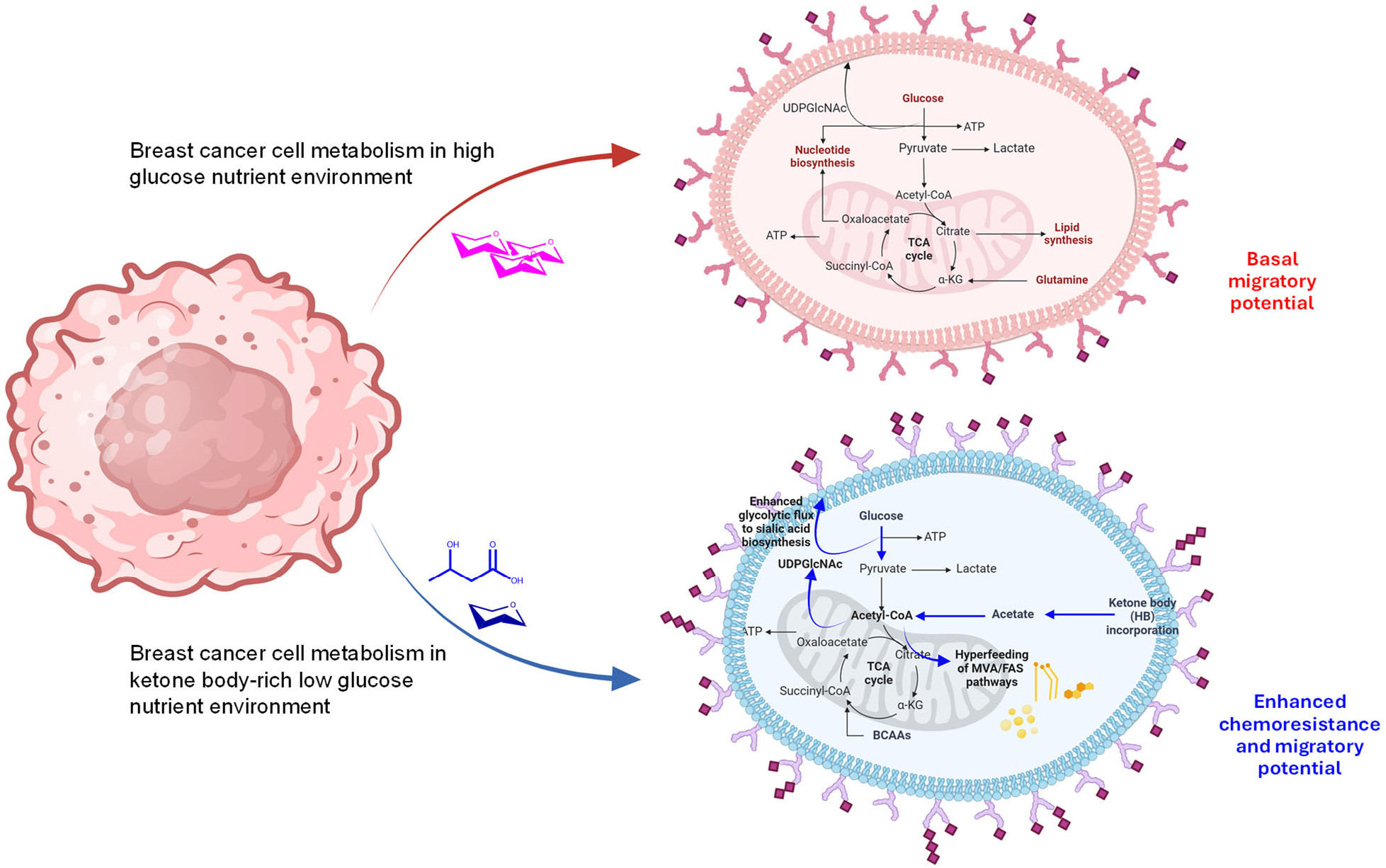
Schematic representation of proposed mechanism highlighting metabolic pathways participating in chemoresistance and invasion in response to ketomimetic nutrient medium. Bold blue arrows indicate the increased turn-over of the specific metabolic pathways. Thin black arrows represent basal-level reaction activity.

## CONCLUSION

We report herein, a mechanistic investigation of the impact of ketone bodies as altered nutrient sources on BC cells. The degree of glycocalyx sialylation, as measured using bioorthogonal click-chemistry, increased significantly in an LG and ketone body-supplemented LG medium. This increase was accompanied by a chemoprotective effect in these cells. Surprisingly, we observed an increase in drug-induced cytotoxicity in the case of non-cancerous mammary epithelial cells and significantly lowered sialylation levels as the media glucose was lowered and HB was added. The differential pattern in cancer vs. normal cells was traced to its biochemical roots, i.e., the upregulation of HBP and lipid accumulation. FLIM measurements indicated that ketone bodies promote a preference for glycolytic metabolic signature. While DOX tends to induce an initial preference for glycolysis in invasive breast cancer cells, a ketomimetic nutrient medium suppresses any further DOX-induced enhancement of glycolysis, correlating with an observed protective mechanism against the chemotherapeutic. Slower proliferation rates were observed for the ketomimetic medium, suggesting a mechanism that BC cells use to escape the antineoplastic effects of a drug like DOX. We find that the glucose taken up by BC cells in a ketogenic medium (via heightened glycolysis) is shunted to the sialic acid biosynthesis that is further supported by production of UDP-GlcNAc from the additional Acetyl CoA parallelly produced from ketone bodies. Chemotaxis studies revealed that ketone bodies promote migratory aptitude of BC cells and in turn enhance their metastatic potential - coincident with enhancement in sialylation.

Measurement of cellular internalization of DOX revealed that the relative changes scale analogous to the toxicity trends across the nutrient media in MCF-10A cells. In the case of the hypersialylated BC cells, there is an increase in internalized DOX upon removal of sialic acids, pointing to the ability of negatively charged sialic acid residues to “capture the DOX at the cell surface” via electrostatic interaction. Taken together, our observations suggest that in a ketomimetic nutrient medium, BC cells utilize a dual machinery of sialylation and lipid accumulation to enhance their chemoresistance and invasive potential. Furthermore, the aberrantly sialylated, hyperbranched glycocalyx promotes metastasis by facilitating cellular migration and immune evasion. On the other hand, ROS quenching arising out of fatty acid synthesis/metabolism coupled possibly with cholesterol build-up from MVA pathway together protect the tumor cells against a chemotherapeutic attack. Since the normal cells lack this machinery, they are not able to utilize ketone bodies as nutrients and die of lipotoxicity. Therefore, our data suggests that a ketogenic diet is not safe for BC patients and can lead to more metastatic lesions. Additionally, a ketogenic diet is expected to make cancer cells more invasive and drug-resistant while increasing the propensity of healthy cells in the body to be damaged by a chemotherapeutic agent being administered. Studies like ours that focus on several chemical biology (metabolic and pro-oncogenic) pathways offer an avenue to better understand effects of dietary interventions and can help formulate informed dietary regimens for patients.

## METHOD DETAILS

### Cell culture

Breast cancer cell lines MCF7 and MDA-MB-231 along with mammary epithelial cells, MCF-10A were procured from ATCC and passage numbers P8-P20 were used for all the experiments. All cells were passaged according to standard ATCC recommended protocols and grown in an incubator that was maintained at 37°C, 5% CO_2_ and 99% humidity. A low-glucose variant of each cell line was generated by passaging an inoculum of the P7 in low-glucose medium and then maintaining in that medium through the subsequent passages. For cells seeded in HBLG for an experiment, a seeding inoculum from the low glucose cells was seeded in 3 mM hydroxybutyrate containing low-glucose media (LG) for the requisite period of the experiment. **Table 1** describes the detailed compositions of the culture media.

**Table 1.**
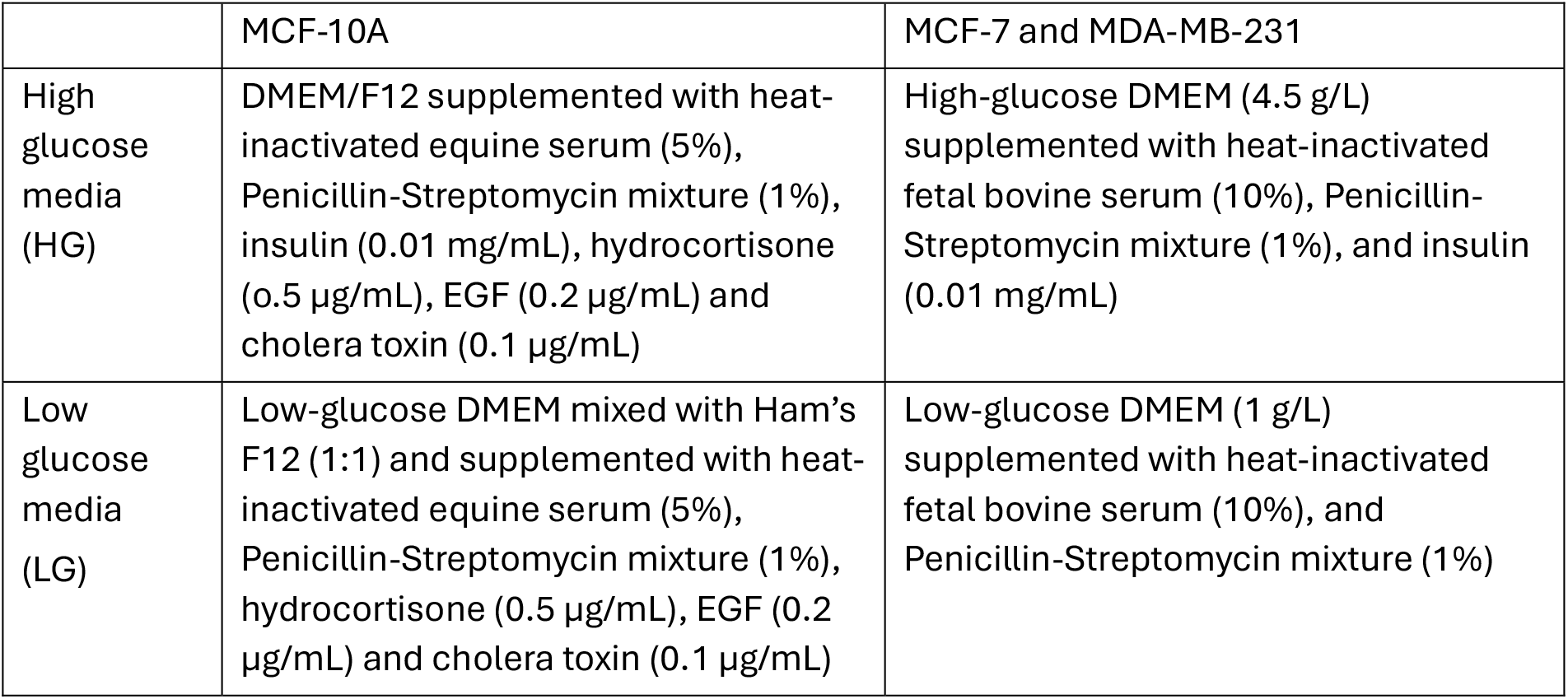
Media compositions used for culturing each cell line prior to seeding for an experiment

|  | MCF-10A | MCF-7 and MDA-MB-231 |
| --- | --- | --- |
| High glucose media (HG) | DMEM/F12 supplemented with heat-inactivated equine serum (5%), Penicillin-Streptomycin mixture (1%), insulin (0.01 mg/mL), hydrocortisone (0.5 µg/mL), EGF (0.2 µg/mL) and cholera toxin (0.1 µg/mL) | High-glucose DMEM (4.5 g/L) supplemented with heat-inactivated fetal bovine serum (10%), Penicillin-Streptomycin mixture (1%), and insulin (0.01 mg/mL) |
| Low glucose media (LG) | Low-glucose DMEM mixed with Ham's F12 (1:1) and supplemented with heat-inactivated equine serum (5%), Penicillin-Streptomycin mixture (1%), hydrocortisone (0.5 µg/mL), EGF (0.2 µg/mL) and cholera toxin (0.1 µg/mL) | Low-glucose DMEM (1 g/L) supplemented with heat-inactivated fetal bovine serum (10%), and Penicillin-Streptomycin mixture (1%) |

### Bioorthogonal imaging of sialic acids

For metabolic labeling and biorthogonal imaging, cells were seeded in an 8-well chamber slide at a density of 2.5 ×10^4^ cells/well. 100 µM of Ac_4_MazNAz was added in the respective media (HG, LG and HBLG) and incubated for 72 h with the cells. At the end of 72 h, CellBrite® Steady Membrane Labeling Kit (Biotium, 30108) was used for staining the cell membranes using the manufacturer’s protocol. The media was then removed, and all the wells were washed twice with PBS, following which a 1 µM dilution of Click-IT DIBO-AF488 (Thermo Scientific) in 1% FBS containing PBS was added and incubated with the cells for 1 h at room temperature. Cells were then washed twice with PBS and maintained in 1% FBS containing PBS for the duration of imaging. Images were acquired with an Olympus FV3000 confocal microscope using UPlanSApo 20x/0.75NA Olympus objective (AF488 ex/em: ∼490/525 nm; CellBrite Steady 650 ex/em ∼ 656/676). Each media condition was assayed in two technical replicates and 8-10 fields of view (FOVs) per well were collected at the time of imaging. In one well, no DIFO-AF488 was added and was used for background subtraction. No Ac_4_ManNAz controls were also tested in initial experiments to confirm specificity of click reaction. Image analysis was performed using a customized pipeline in Cell Profiler that used the images in the membrane stain channel to generate a binary and (a) calculate the total perimeter of cell membrane in the FOV, and (b) create a mask on the images from the AF488 channel which was then used to compute total intensity of click reacted-azidosialic acids on the cell membranes in that FOV. Intensity/length of perimeter was then computed for each condition and plotted relative to the high-glucose condition using GraphPad Prism.

### LDH Cytotoxicity Assay

For determination of drug response, cells were seeded in the respective media (HG, LG and HBLG) in the wells of a 96-well plate at a density of 1 × 10^4^ cells/well. After 24 h, 1 µM solution of DOX (in the respective media) was added to one set of wells and incubated for another 48 h. Control wells received plain media, without DOX. CyQUANT™ LDH Cytotoxicity Assay, fluorescence (Thermo Scientific C20302) was then performed using the manufacturer’s protocol and measurements were conducted on a Cytation 5 (BioTek) plate reader. The extent of cytotoxicity was computed relative to the untreated controls and values were then normalized against the high glucose condition for comparison, when plotted using GraphPad Prism.

### Fluorescence Lifetime Imaging Microscopy (FLIM)

Cells were seeded at a density of 2.5 × 10^5^ cells/well in 250 µL of the respective media (HG, LG and HBLG) on the surface of one well of three separate 8-well chamber glass slides (one slide/sample). After three days, the media was replenished with 250 µL of phenol red free media of the same composition as before and set on the heated stage (maintained at 37 °C) of the FLIM microscope to begin imaging. After taking the images for t = 0 for 3 fields of view, 5 µL of DOX solution (50 µM) was added to the well and gently mixed to minimize disturbance to the layer of cells. Images of the same three fields of view were then captured every 15 min for a total of 60 min from the time of starting. The fluorescence lifetime images were obtained from the FLIM system (Alba v5, ISS) which was equipped with the Nikon inverted microscope with 60x, NA 1.1 water objective (NALUMFLN60XW, Olympus) and the 405 nm diode laser. The laser line passed a multi-band dichroic mirror to excite the samples. The fluorescence emission light traveled back along the same path to the multi-band dichroic mirror, and finally went into the avalanche photodiode detector (SPCM-AQR-15, Perkin Elmer) with 200 nm pinhole and corresponding bandpass filter (445/40nm, Semrock) for NADH measurement. The current setup was also equipped with an ASI XY automatic stage with motorized Z control. The software (VistaVision) we use can record the position of different FOV. This allows us to trace back the same cells after the DOX was added. The photon counts were acquired with the FastFLIM unit to build up the phase histogram. To calibrate the FLIM system, we used a fluorescence lifetime standard, Atto 425, with a reference lifetime of 3.6 ns, before imaging samples in this experiment. The laser repetition period was 50 ns, and it was divided into 256 bins. FLIM images were scanned 5 times with dwell time of 0.2 ms/pixel. The data were collected by the confocal scanning system from a 256 × 256-pixel area (100 × 100 μm field of view) before and after DOX treatment.

### FLIM Data analysis

The FLIM data was analyzed in digital frequency-domain (DFD) by taking the Fourier transform of the fluorescence decay histogram. In other words, the fluorescence decay histogram of each pixel was presented as a single point in the phasor plot under corresponding modulation frequency condition. The modulation ratio and phase shift were then extracted from the point in the phasor plot. With the modulation ratio and phase shift data over multiple modulation frequencies, they were fitted to the fluorescence decay model to obtain the fluorescence lifetime of interest. The fluorescence lifetime standards fit to the one-component decay model during the calibration stage. In living (cancer) cells, we expect there are two populations contributing to the fluorescence decay measurements at each pixel: free and protein-bound NADH. Fixing the 0.3 ns as one lifetime component for free NADH (*τ*_*1*_), we analyzed the fluorescence lifetime measurements (*m(t)*) in cancer cells with bi-exponential decay model,

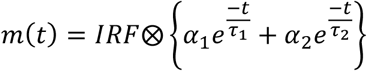

Where *α*_*1*_ is fraction amplitude of free NADH, *α*_*2*_ and *τ*_*2*_ are fraction amplitude and fluorescence lifetime of protein-bound NADH, and *IRF* can be obtained from the experiment. By definition, we then determined the NADH metabolic index by NAD+/NADH ratio,

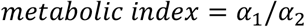

### Glucose Consumption Assay

Cells were seeded in a 6-well plate at a density of 1 × 10^5^ cells/well in 2 mL of the respective media (HG, LG and HBLG). In parallel, 2 mL of the same media was added to another set of wells (without cells) for control and the plate was incubated at 37 °C, 5% CO_2_ and 99% humidity for 4 days. After this, the supernatant media was collected from the wells and spun at 3000 rpm for 5 min at 4 °C to remove any cells/cell debris. The clarified media was used for assessment of glucose concentration using Glucose Colorimetric Assay Kit from Cayman Chemicals (Item No. 10009582). Glucose standards provided by the manufacturer were used for obtaining calibration curve and sample supernatants were used at 1:40 dilution in the assay buffer (1:80 for high glucose media). The amount of glucose consumed by the cells was obtained by subtracting the values of glucose concentration in media incubated in the wells without cells from those that were incubated with cells. All measurements were conducted on a Cytation 5 plate reader (Biotek) following manufacturer’s protocol. The total amounts of glucose consumed in each nutrient condition were plotted using GraphPad Prism.

### Cellular Proliferation Measurement

In a 48-well plate, cells were seeded at a density of 2 × 10^3^ cells/well in 500 µL medium, with duplicates for each day and nutrient condition (HG, LG and HBLG). On the fourth day following seeding (day zero), cells were trypsinized from the respective wells and counted using Countess after Erythromycin B staining every 24 h. Media in the wells was replaced every three days for the cells maintained for longer durations. Also, depending on cell-type, cells were transferred to 24/12-well/6-well plates as and when confluency reached 80%. Each counting was repeated twice, and total average cell counts were plotted against day points using GraphPad Prism to obtain proliferation curves.

### Cellular Migration Assessment

Corning transwell inserts (Sigma, CLS3422) were used for assessment of cellular migratory aptitudes. Initially, cells were seeded in a 6-well plate at a density of 1 × 10^5^ cells/well in 2 mL of the respective nutrient media (HG, LG and HBLG). After three days, the media was replaced with 1 mL of 0.5% FBS containing media of the same kind (only FBS amount was altered). After a 24 h incubation period, cells were trypsinized and seeded onto transwell inserts (in 250 µL of 0.5% FBS containing media) at a density of 1.5 × 10^3^ cells/well. The transwell inserts were gently placed in the wells of a 24-well plate that were filled with 500 µL of 20% FBS containing media of the same kind (HG, LG or HBLG) to provide chemotactic stimulus. After 24 h, the transwell inserts were recovered carefully, the upper surface was wiped with wet cotton swabs and gently washed with PBS. This was followed by fixation using ethanol, more PBS washes of the upper surface and staining the cells on the lower surface using crystal violet (0.2%). The rear (lower) sides of the inserts were imaged under bright field using UPlanSApo 20x/0.75NA Olympus objective and color CMOS camera (Thorlabs, CS165CU) on a home-built microscope (Thorlabs EDU OMC1). The percentage of total area occupied by cells was computed using ImageJ taking at least 10 FOVs per transwell in an experiment. For statistical stringency, the whole experiment was repeated twice, and values pooled to obtain the final graph.

### Real-time quantitative PCR for gene expression analysis

Different cell types grown in their basal media (HG) were harvested and seeded in 90 mm culture dishes at a density of 5 × 10^5^ cells/dish. When cells reached ∼70% confluence they were washed with PBS quickly, scraped off using 1 mL of RNA lysis buffer (part of RNA prep kit) and frozen at -80 °C. After at least 30 min-1 h, the lysates were thawed on ice and RNA was extracted using *Quick*-RNA Miniprep Kit (Zymo Research). RNA concentration was measured using nanodrop and then 1 µg of total RNA from each sample was taken for cDNA synthesis using High-Capacity cDNA Reverse Transcription Kit (Applied Biosystems). cDNA was diluted 1:10 and used for qPCR reactions employing Luna® Universal qPCR Master Mix (New England Biolabs). For all genes, 1 µL of the cDNA dilution was used per 20 µL reaction and for *FASN*, 2 µL of the cDNA dilution was used (due to low copy number and undetectable Ct values when using 1 µL). Ct values were computed by the instrument and used for calculation of ΔΔCt using *ACTB* (Beta-actin) as a housekeeping control. The table of primers used in the study is included in **Table S1**. Experiments were performed with three independent biological replicates each having at least two (or more) technical replicates.

### Cellular internalization of DOX

DOX fluorescence was used to assess the amount of drug that has internalized in the cells in response to removal of sialic acid residues from the glycocalyx. Since the action of sialidase enzyme occurs in slightly acidic pH, close to that of RPMI, cells used for this experiment were separately grown in media formulations where RPMI was used in place of DMEM (HG, LG and HBLG). No glucose RPMI was purchased from Thermo Scientific (catalog no. 11879020) and used to prepare the high and low glucose containing media to resemble the low (1g/L) and high glucose (4.5 g/L) DMEM. Three days before the experiment, 2 × 10^4^ cells/well were seeded in an 8-well chamber slide in 250 µL media/well. Two days later, media in half of the wells was replaced with serum free media of the same kind (HG, LG, or HBLG RPMI) containing sialidase enzyme (Millipore Sigma 10269611001) at a dilution of 1: 100 and incubated overnight at 37 °C, 5% CO2. The control wells of each media kind did not receive the enzyme. The cells were then treated with a 1 µM dose of DOX for 1 h after which they were washed and stained with DRAQ5 for nuclear counterstain. Imaging was performed with an Olympus FV3000 confocal microscope using UPlanSApo 20x/0.75NA Olympus objective. Images of at least 10 FOV’s were acquired per well. One well did not receive DOX treatment and was used for background subtraction for DOX fluorescence intensity (I_blank_). Image analysis was performed using customized pipeline in Cell Profiler where images of the DRAQ5 channel were used for counting the number of cells in each FOV and DOX fluorescence intensity (after background subtraction) was divided by number of cells for the respective FOV. The formula for each FOV thus reads as:

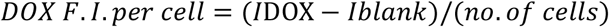

Fold change in internalized DOX was calculated relative to cells in the corresponding nutrient condition without sialidase and plotted using GraphPad Prism. For confirmation of cleavage of sialic acids in the cells, bioorthogonally click-labeled sialic acid channel was also imaged in a few of the repeat runs using 100 µM ManNAz in the media and 5 µM of DIBO-AF 647 at the time of click reaction. For these set of experiments, DAPI was used as a nuclear counterstain and imaging was performed in a Nikon CSU-W1 Spinning Disk Confocal System using Nikon Plan Apo λ 20x/0.75 objective

### Assessment of Lipid Droplet accumulation

BODIPY staining was used for imaging of lipid droplets inside the cells. On day zero, cells were seeded in an 8-well chamber slide at a density of 2.5 × 10^4^ cells/well in 250 µL of the respective media (HG, LG and HBLG). On day 3, the media was aspirated, cells were washed twice with PBS and incubated with 2 µM BODIPY along with 5 µM DRAQ5 at room temperature for 15 minutes. Following two washes with PBS again, the cells were imaged under Olympus FV3000 confocal microscope with a UPlanSApo 20x/0.75NA Olympus objective at 2X digital zoom (BODIPY ex/em: ∼505/515 nm; DRAQ5 ex/em ∼ 633/695 nm) taking at least 10-15 FOV’s per condition. One well did not receive BODIPY, and its avg. intensity used for background subtraction (I_blank_). Image analysis was performed using a customized pipeline in Cell Profiler where images taken in the DRAQ5 channel were used for counting the number of cells in each FOV and BODIPY fluorescence intensity (after background subtraction) was divided by number of cells for the respective FOV. The formula for each FOV thus reads as:

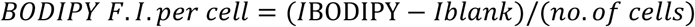

Fold change in BODIPY F.I. per cell was calculated relative to cells grown in HG medium and plotted using GraphPad Prism.

### MALDI-Mass Spectrometry Imaging

For identification of metabolomic profiles of cells fed on different nutrient media, MALDI based mass spectrometric imaging was performed. First, an equal number of cells were seeded on 90 mm dishes in the respective nutrient media (HG, LG and HBLG) and grown for three days at 37 °C, 5% CO_2._ On day 3, nearly 1 million cells from each dish were collected by centrifugation at 120g for 7 minutes. The cells were then washed twice with 150 mM of ammonium acetate buffer and cell pellets (with negligible liquid on them) were frozen in -80 °C. At the time of imaging, cell pellets were sectioned at 12 μm thickness and sections were coated with 7 mg/mL N-(1-naphthyl) ethylenediamine dihydrochloride in 70% methanol over 8 passes using an HTX M5 Robotic Reagent Sprayer (HTX Technologies, Chapel Hill, NC). Spray conditions were as follows: a nozzle temperature of 75°C, a nozzle height of 40 mm, a track speed of 1200 mm/min, a track spacing of 2 mm, a flow rate of 0.12 mL/min, and a nitrogen pressure of 10 psi. Sections were imaged at 50 μm resolution in negative ion mode using a Bruker timsTOF fleX QTOf mass spectrometer (Bruker Daltonics, Billerica, MA). Instrument acquisition parameters were as follow: *m/z* range 50-1000, 500 shots per pixel, a Funnel 1 RF of 125.0 Vpp, a Funnel 2 RF of 150.0 Vpp, a Multipole RF of 175.0, a Collision Energy of 5.0 eV, a Collision RF of 500.0 Vpp, a Transfer Time of 65.0 μs, and a Pre Pulse Storage of 4.0 μs. Matrix was removed after imaging and sections were stained with H&E for visualization of the cells. MS spectra were compiled into a single image file with each peak from the average spectrum being displayed as a function of its spatial position and relative intensity across the section. Images were visualized using SCiLS Lab Pro 2024b and putative IDs were generated using MetaboScape followed by manual search for missing ID’s using Metaspace2020. Normalized root mean square intensities were then compared across all conditions by performing a pixel-wise Principal Component Analysis using the SCiLS Lab software. Heat maps for the enrichment analysis were obtained using Heatmapper.com.^43^ For cholesterol assesment, another set of sections from the same set of samples were imaged at 50 μm resolution in positive ion mode.

### Statistical Analyses

All methods and software used for statistical analysis have been mentioned in the respective figure legends and method details for each experiment. The following details have been included in the figure legends: the statistical tests used, exact value of n (technical replicates), exact value of N (biological replicates), definition of symbols, error bars and precision measures (i.e. mean, median, SD, and SEM as the case may be). Significance was defined using the respective p values as follows: ns = p > 0.05, * = p < 0.05, ** = p < 0.01, and *** = p < 0.001. For imaging experiments, the no. of fields of view per condition have also been elucidated in the respective section of method details.

## ASSOCIATED CONTENT

### Supplemental Information

Document S1. Figures S1–S7 and Table S1: This document includes additional images and supporting data analysis, and data from control studies for some experiments along with the table of primers used for rt-*q*PCR. For any raw data/image analysis pipelines, please contact the corresponding author, all data is available upon request.

## Author Contributions

M.K. and S.H.P. conceived the idea. M.K. designed the experiments, performed most of the experiments and wrote the initial manuscript draft. Y-IC. performed the FLIM imaging, post-processing and image analysis while also contributing to manuscript writing and data interpretation along with H-CY. P.D. assisted M.K. in cell culture. E.S. performed the MALDI-MSI studies at the Mass Spectrometry Imaging Facility, UT Austin. S.K.S., A.B., and S.H.P. supervised the study and procured resources. M.K. and S.H.P. wrote the manuscript with input and comments from all authors.

## ACKNOWLEDGMENTS

We are grateful to the Mass Spectrometry Imaging Facility at UT Austin funded by Cancer Prevention and Research Institute of Texas (CPRIT) RP190617 award. We sincerely thank Christian M. Jennings for helping with the setup of Thorlabs EDU-OMC1 microscopy kit and Thomas E. Yankeelov for allowing access to the BioTek Cytation 5 instrument that was purchased with a CPRIT RR160005 award. We also thank the Center for Biomedical Research Support Microscopy and Imaging Facility at UT Austin (RRID:SCR_021756) where the spinning disk confocal microscopy was performed. Artwork used for generating the schematic in Figure 6 was prepared using resources from ChemBioDraw 22.2.0 and Biorender.com.

