## Supplemental Data and Tables for "Ketomimetic Nutrients Trigger a Dual Metabolic Defense in Breast Cancer Cells"

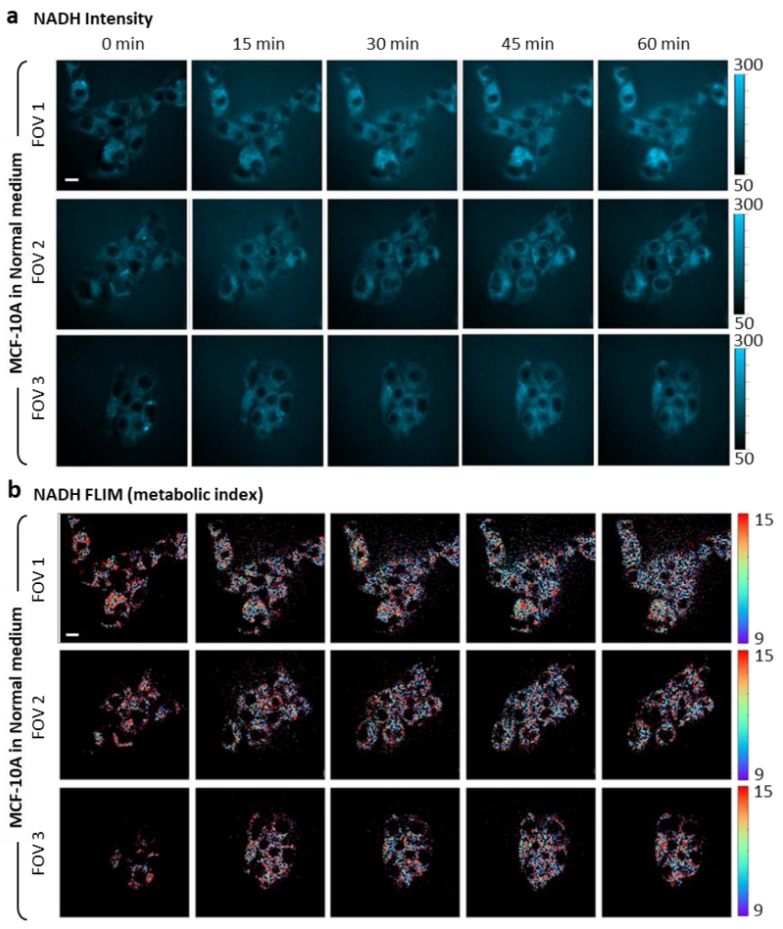


**Supplementary Figure S1.** NADH intensity (a) and metabolic index (b) of MCF-10A cells in normal medium with increasing time of DOX treatment. Three field of views (FOV) taken. Scale bar = 10 μm.

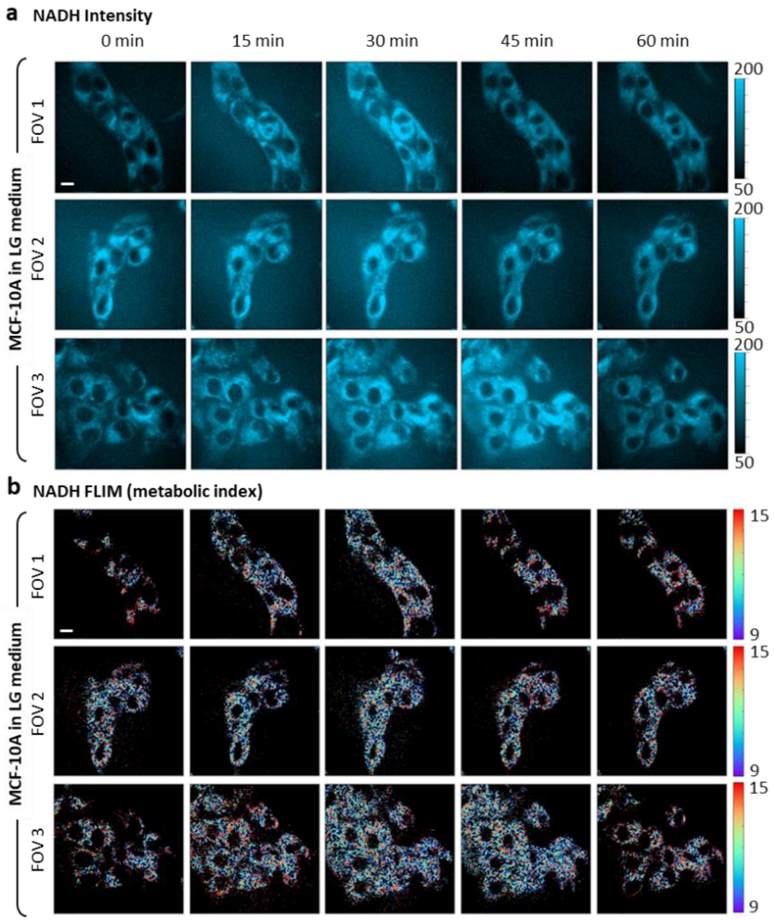


**Supplementary Figure S2.** NADH intensity (a) and metabolic index (b) of MCF-10A cells in LG medium with increasing time of DOX treatment. Three field of views (FOV) taken. Scale bar = 10 μm.

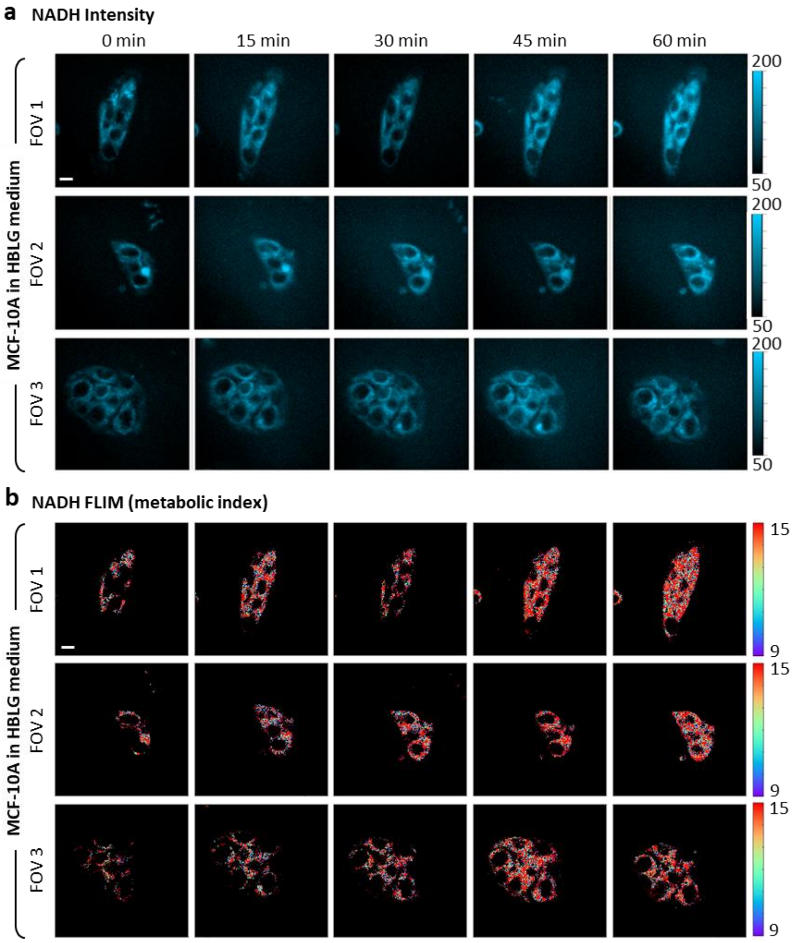


**Supplementary Figure S3.** NADH intensity (a) and metabolic index (b) of MCF-10A cells in HBLG medium with increasing time of DOX treatment. Three field of views (FOV) taken. Scale bar = 10 μm.

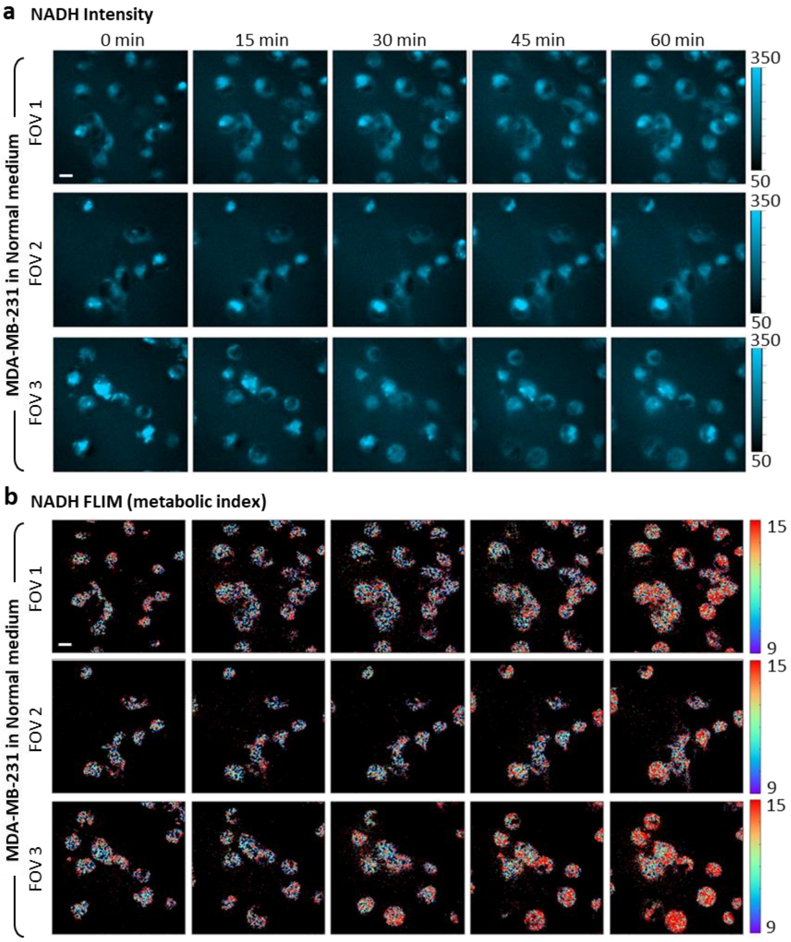


**Supplementary Figure S4.** NADH intensity (a) and metabolic index (b) of MDA-MB-231 cells in normal (HG) medium with increasing time of DOX treatment. Three field of views (FOV) taken. Scale bar = 10 μm.

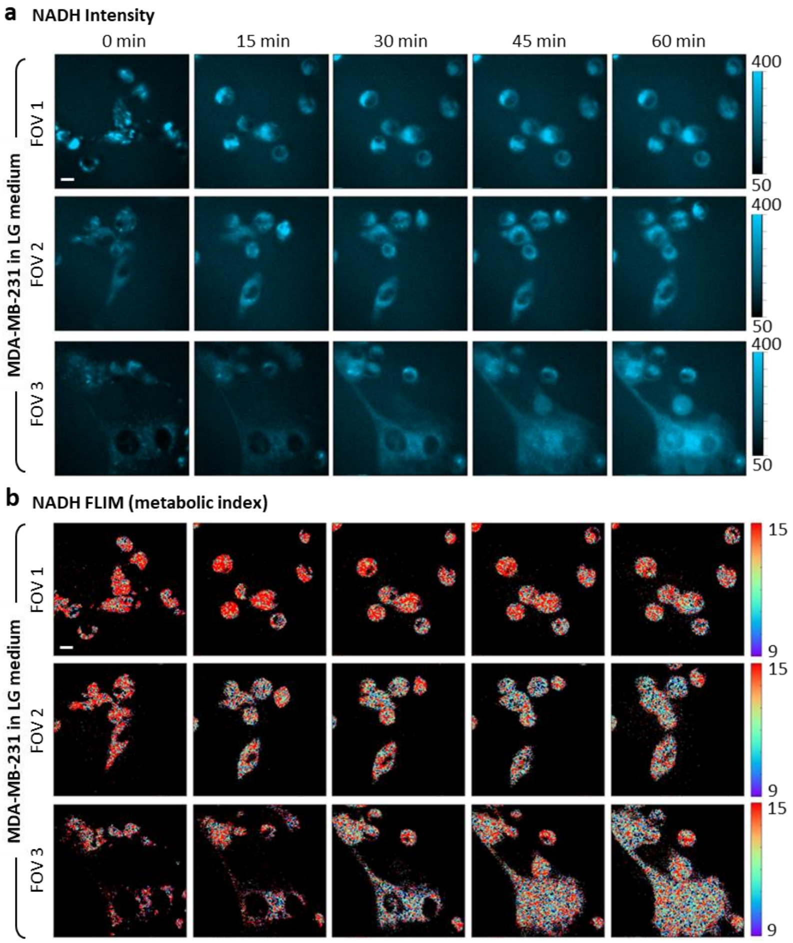


**Supplementary Figure S5.** NADH intensity (a) and metabolic index (b) of MDA-MB-231 cells in LG medium with increasing time of DOX treatment. Three field of views (FOV) taken. Scale bar = 10 μm.

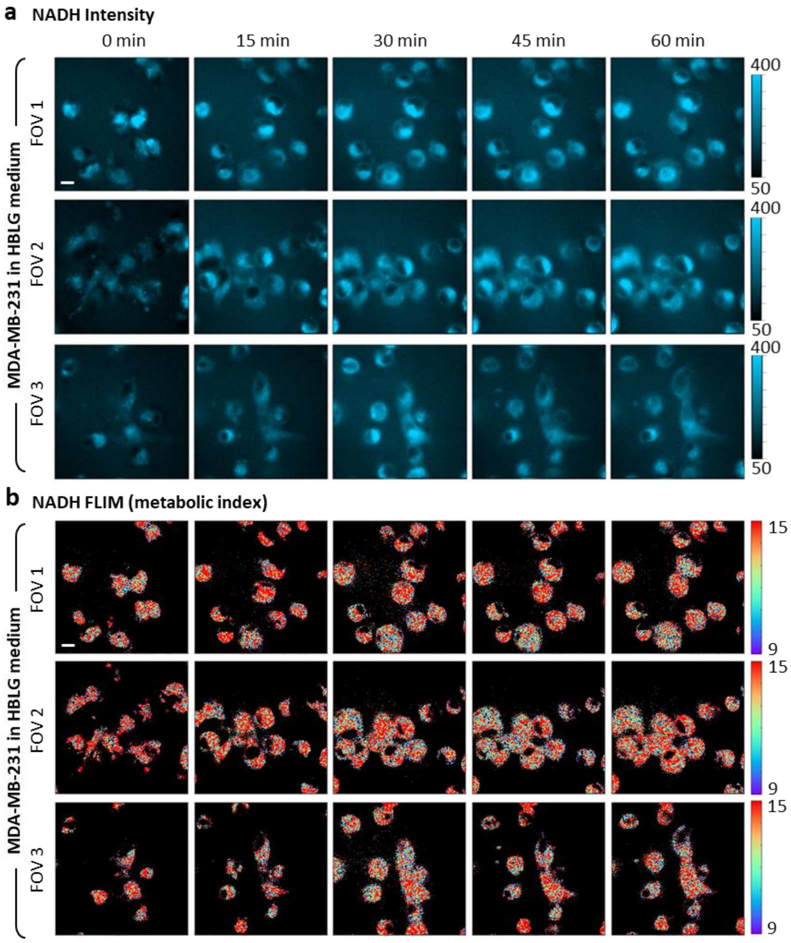


**Supplementary Figure S6.** NADH intensity (a) and metabolic index (b) of MDA-MB-231 cells in HBLG medium with increasing time of DOX treatment. Three field of views (FOV) taken. Scale bar = 10 µm.

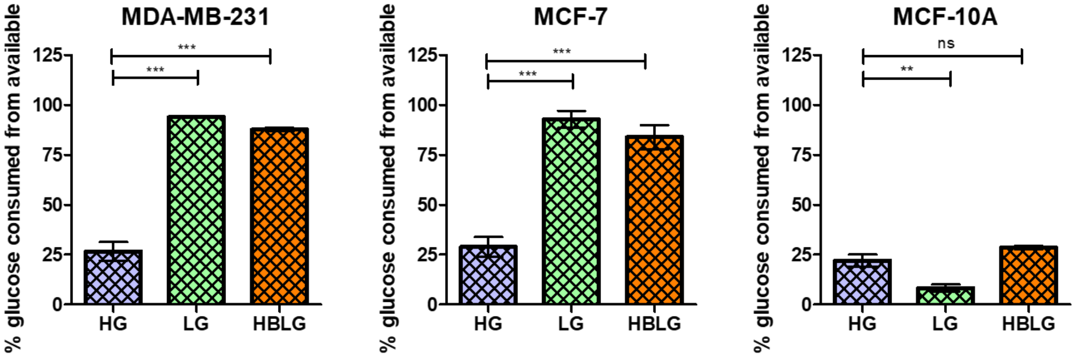


**Supplementary Figure S7.** Glucose consumption patterns (in % of available glucose in the media) of MDA-MB-231, MCF-7 and MCF-10A cells in the respective nutrient media: high glucose (HG), low glucose (LG) and HBLG (hydroxybutyrate containing low glucose). Enzymatic assay based colorimetric detection was used and changes in the amount of glucose consumed by the cells in 4 days was plotted with data from n=3 and N=2 being pooled together. GraphPad Prism was used for plotting the data and unpaired t-tests were performed for column-wise statistical analysis. *, **, *** indicate $p<0.05$, $p<0.01$, and $p<0.001$, respectively.

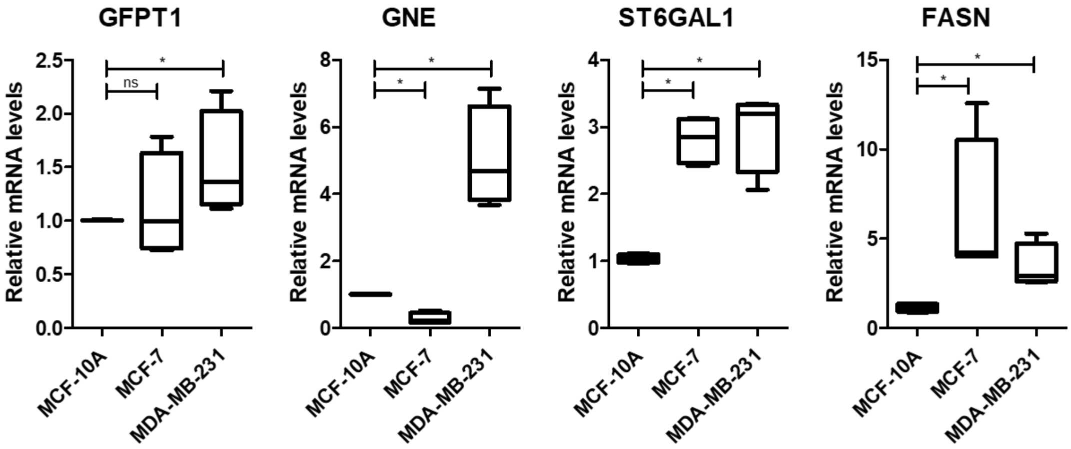


**Supplementary Figure S8**. Box plots representing relative normalized RNA expression of specific enzymes involved in HBP, sialylation and fatty acid synthesis across the cell lines used in the study. Lines are drawn at the median values, with box edges showing the upper and quartiles and whiskers indicating min/max. Mann-Whitney unpaired two-tailed t-test was performed in GraphPad Prism to assess significance, ns and * indicate $p>0.05$ and $p<0.05$, respectively.

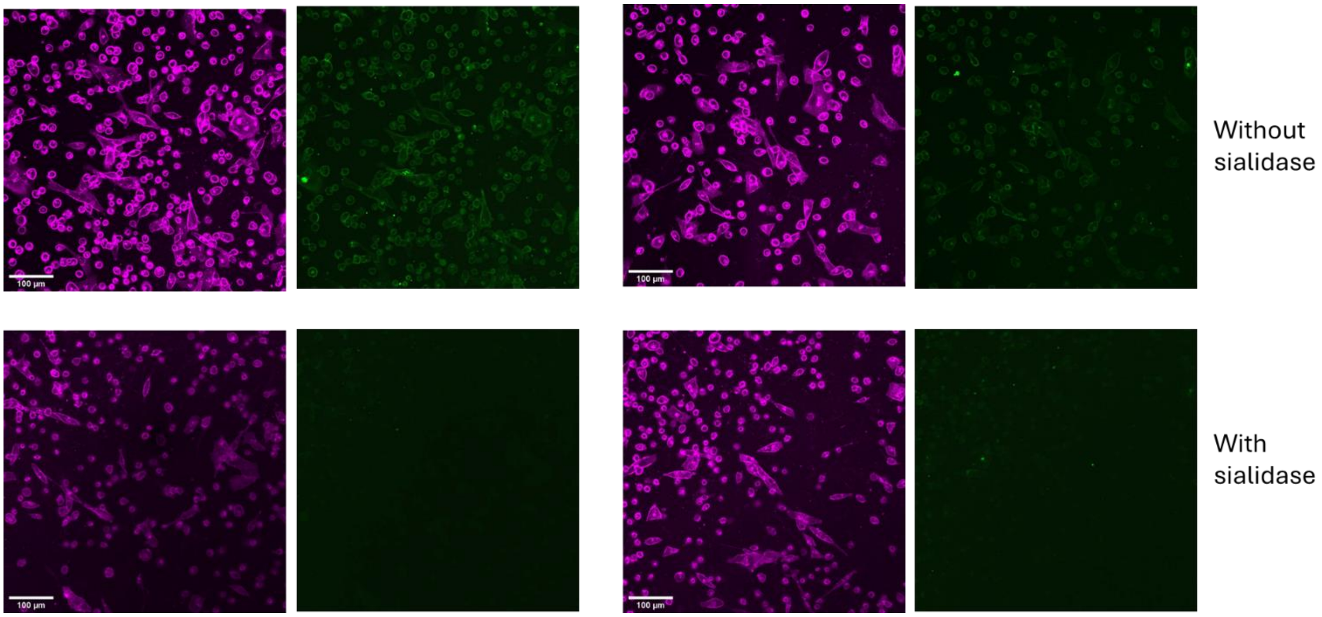


**Supplementary Figure S9.** Representative DIBO AF488 images of MDA-MB-231 cells with and without an overnight treatment of sialidase (1:100) in RPMI. There are two representative field of views for each condition and the membrane labeling channel is shown alongside on the left of each respective AF488 channel. Images were captured using a Nikon CSU-W1 Spinning Disk Confocal System (20X air objective, NA= 0.75). Imaging was performed with an Olympus FV3000 confocal microscope using UPlanSApo 20x/0.75NA Olympus objective. AF488 ex/em: ~490/525 nm; CellBrite Steady 650 ex/em ~ 656/676. All images were collected at identical conditions and are displayed with the same LUT.

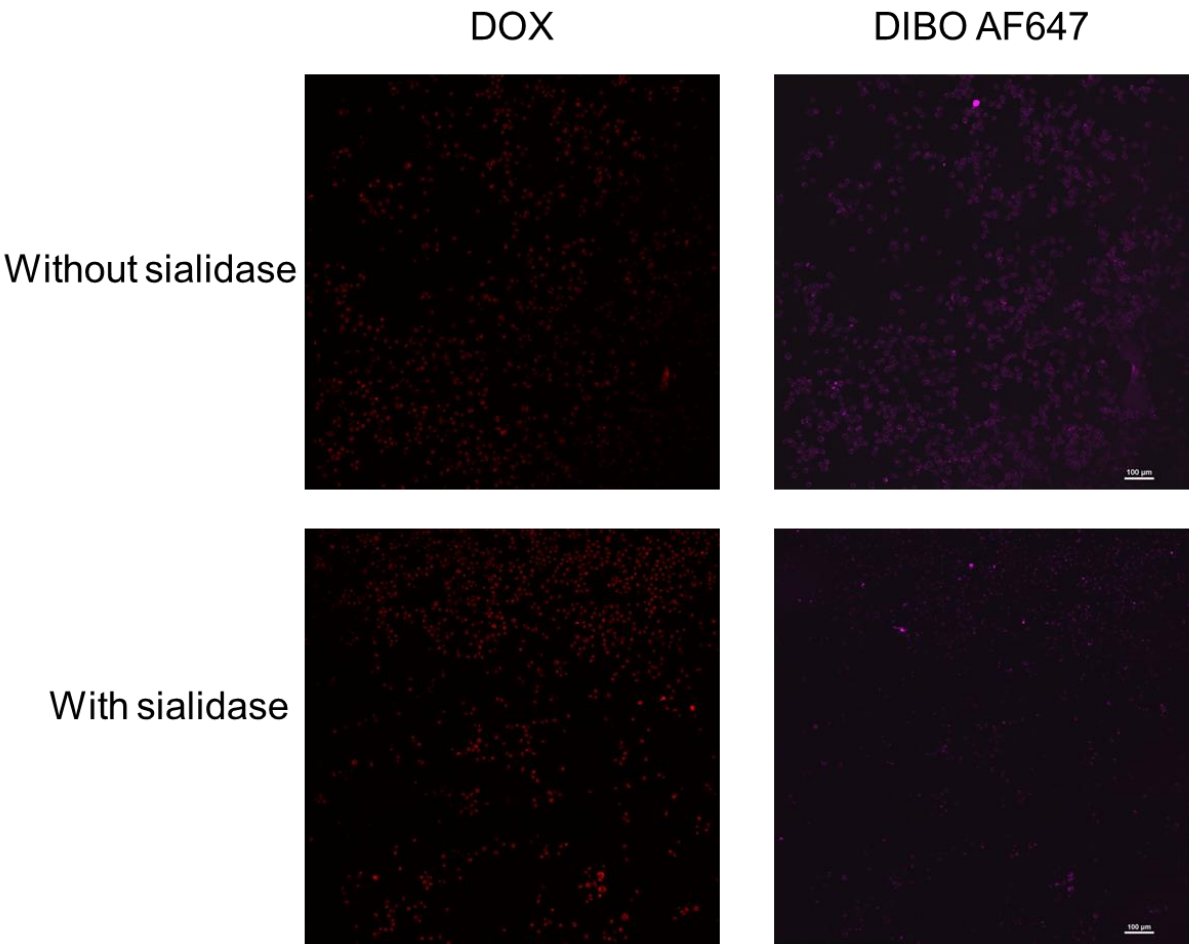


**Supplementary Figure S10.** Tile scan (3X3) images of MDA-MB-231 cells treated with 1uM DOX for 1 h with and without an overnight sialidase incubation. Images were captured using a Nikon CSU-W1 Spinning Disk Confocal System (20X air objective, NA= 0.75). DAPI ex/em: ~405/460 nm; DOX ex/em ~488/561 nm; AF647 ex/em~ 650/670). All images were collected at identical conditions and are displayed with the same LUT.

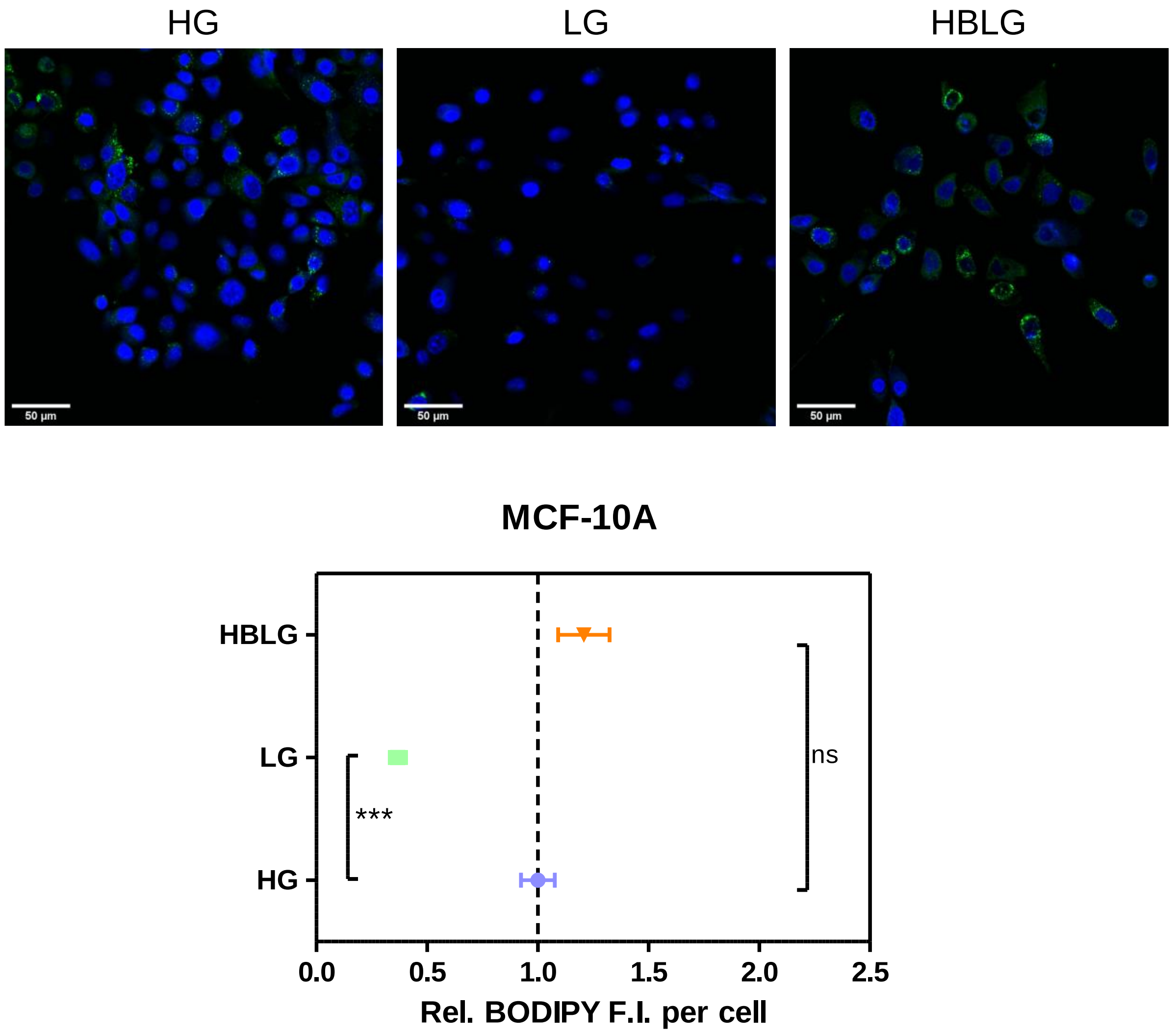


**Supplementary Figure S11.** Lipid droplet imaging in MCF-10A, where the upper panel shows Representative images of MCF10A cells obtained using BODIPY staining for detection of lipid droplets in the respective nutrient media. DRAQ5 was used as a nuclear counterstain for cell count. Green: BODIPY 505, Blue: DRAQ5; scale bar: 50 µm. All images were collected at identical conditions and are displayed with the same LUT. Lower panel shows the graph obtained from quantification of images obtained from four biological replicates of the experiment with at least 10 different FOVs collected for each experiment. ns and *** indicate $p > 0.05$ and $p < 0.001$, respectively.

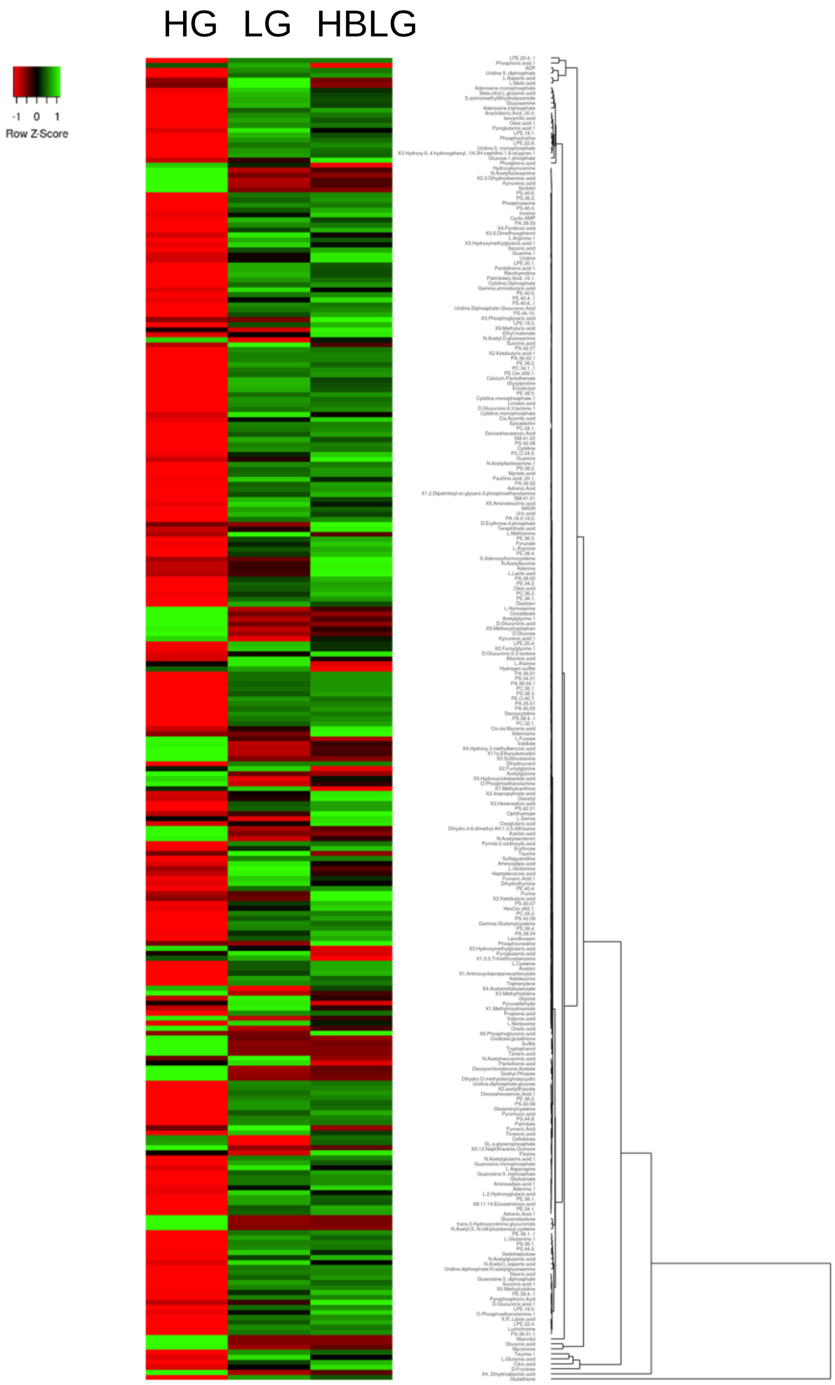


**Supplementary Figure S12.** Heatmap and dendrogram analysis of metabolites identified (in negative-ion mode MALDI-MSI) in MDA-MB-231 cells grown in different nutrient media, HG, LG and HBLG. (Clustering Method: Average Linkage; Distance Measurement Method: Euclidean).

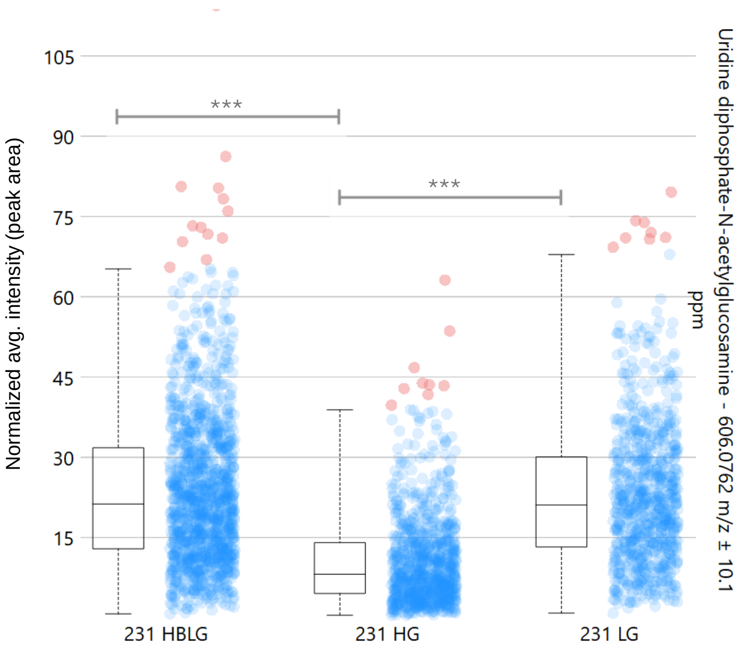


**Supplementary Figure S13.** Relative distributions of Uridine diphosphate-N-acetylglucosamine in MDA-MB-231 cells grown in different nutrient media, HG, LG and HBLG. Root mean square normalization was followed for computing average intensities and Wilcoxon rank sum corrected p values were used for assessing statistical significance where *** indicates p < 0.001.

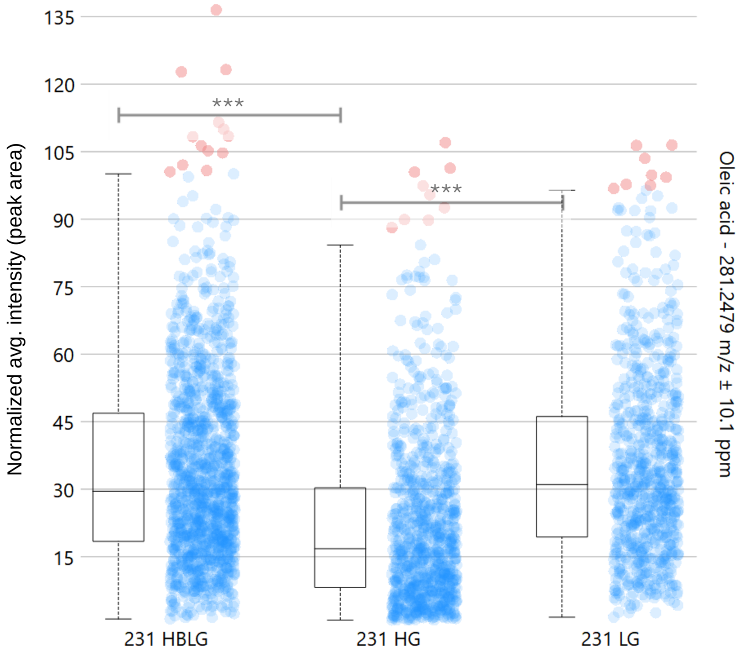


**Supplementary Figure S14.** Relative distributions of Oleic acid in MDA-MB-231 cells grown in different nutrient media, HG, LG and HBLG. Root mean square normalization was followed for computing average intensities and Wilcoxon rank sum corrected p values were used for assessing statistical significance where *** indicates p < 0.001.

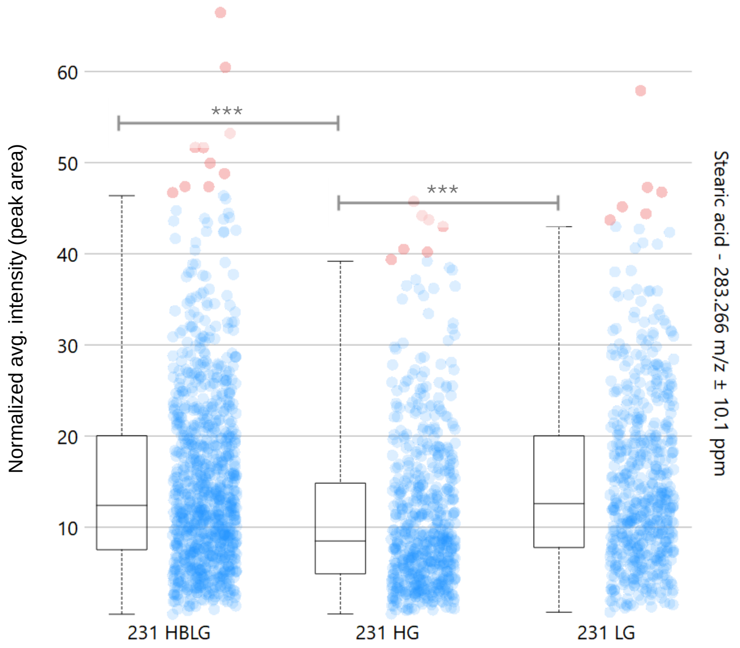


**Supplementary Figure S15.** Relative distributions of Stearic acid in MDA-MB-231 cells grown in different nutrient media, HG, LG and HBLG. Root mean square normalization was followed for computing average intensities and Wilcoxon rank sum corrected p values were used for assessing statistical significance where *** indicates $p < 0.001$.

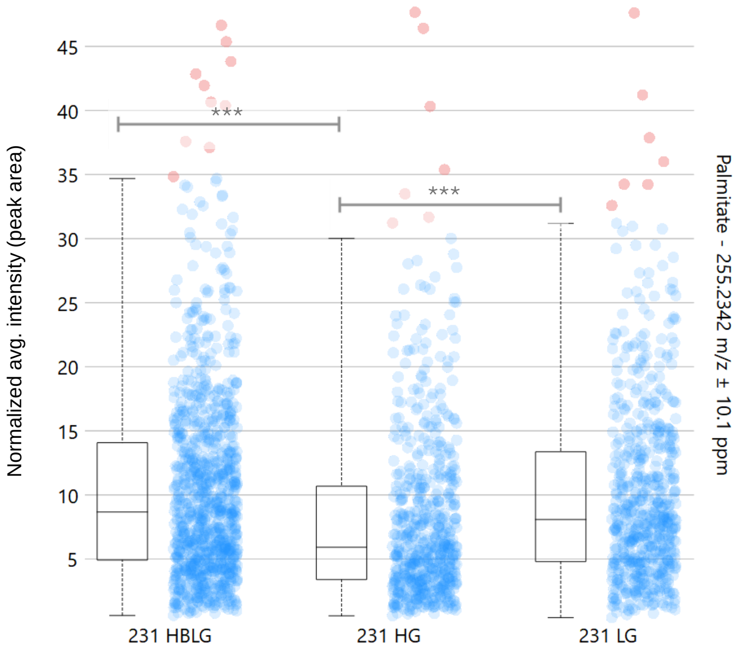


**Supplementary Figure S16.** Relative distributions of Palmitate in MDA-MB-231 cells grown in different nutrient media, HG, LG and HBLG. Root mean square normalization was followed for computing average intensities and Wilcoxon rank sum corrected p values were used for assessing statistical significance where *** indicates $p < 0.001$.

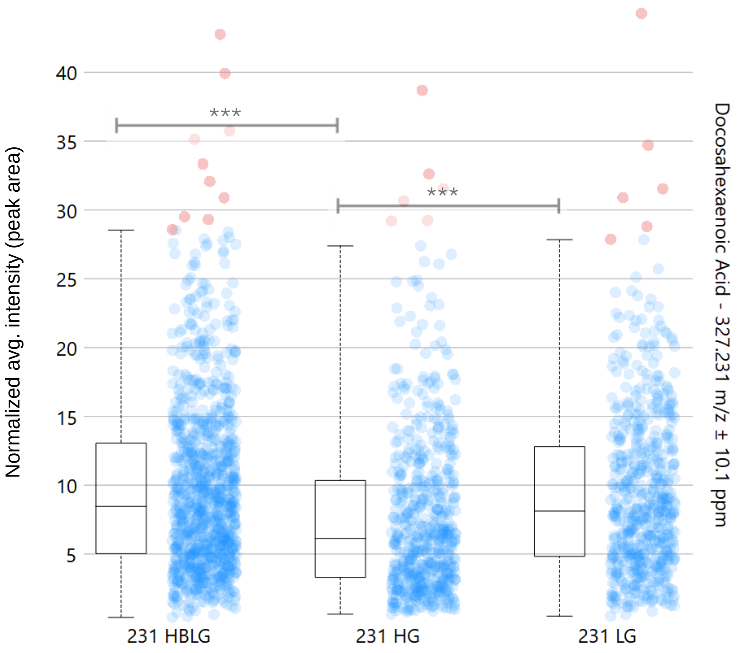


**Supplementary Figure S17.** Relative distributions of Docosahexaenoic acid in MDA-MB-231 cells grown in different nutrient media, HG, LG and HBLG. Root mean square normalization was followed for computing average intensities and Wilcoxon rank sum corrected p values were used for assessing statistical significance where *** indicates p < 0.001.

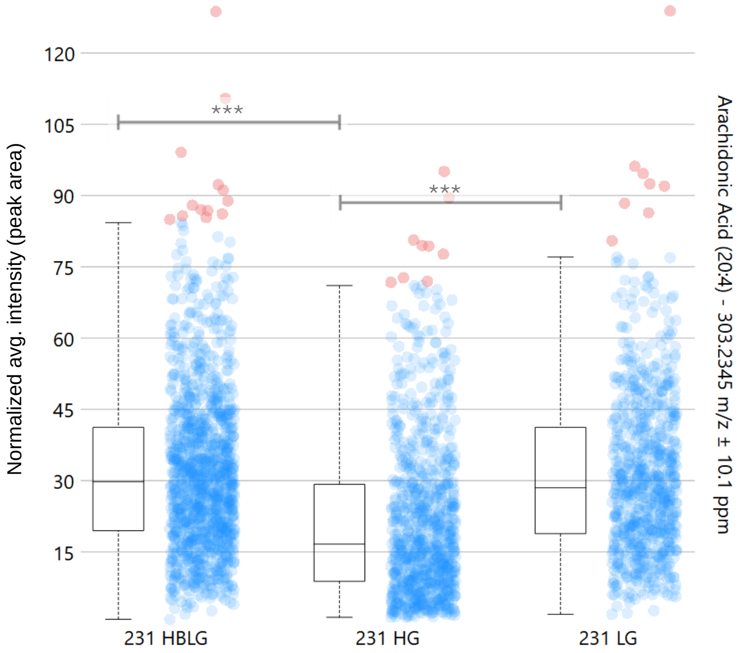


**Supplementary Figure S18.** Relative distributions of Arachidonic acid in MDA-MB-231 cells grown in different nutrient media, HG, LG and HBLG. Root mean square normalization was followed for computing average intensities and Wilcoxon rank sum corrected p values were used for assessing statistical significance where *** indicates p < 0.001.

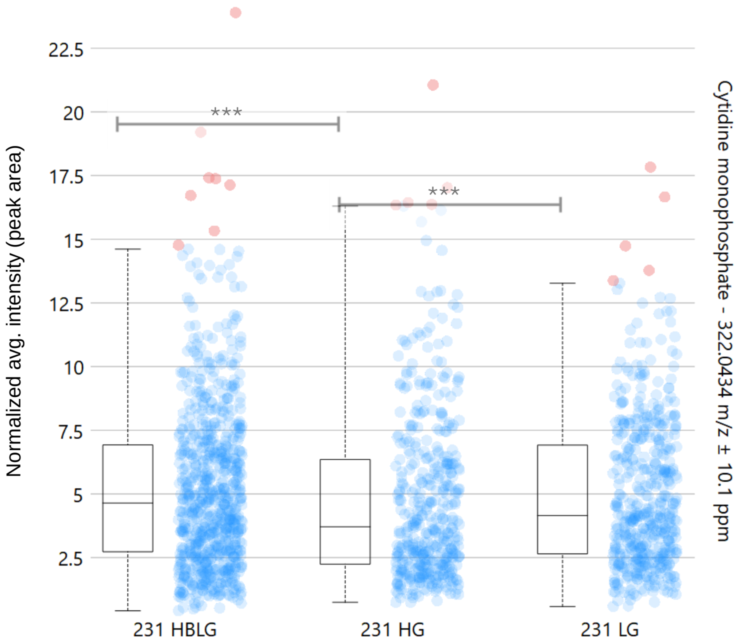


**Supplementary Figure S19.** Relative distributions of Cytidine monophosphate (CMP) in MDA-MB-231 cells grown in different nutrient media, HG, LG and HBLG. Root mean square normalization was followed for computing average intensities and Wilcoxon rank sum corrected p values were used for assessing statistical significance where *** indicates p < 0.001.

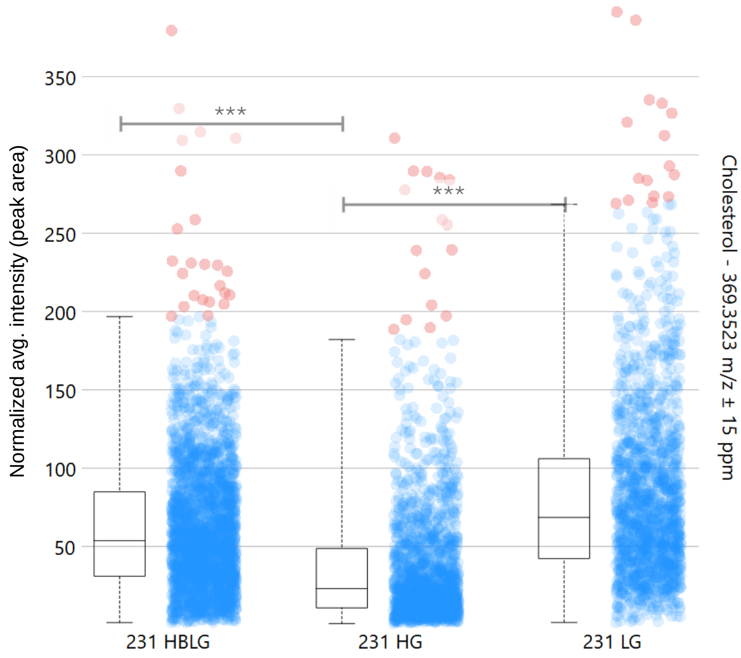


**Supplementary Figure S20.** Relative distributions of Cholesterol in MDA-MB-231 cells grown in different nutrient media, HG, LG and HBLG. Root mean square normalization was followed for computing average intensities and Wilcoxon rank sum corrected p values were used for assessing statistical significance where *** indicates p < 0.001.

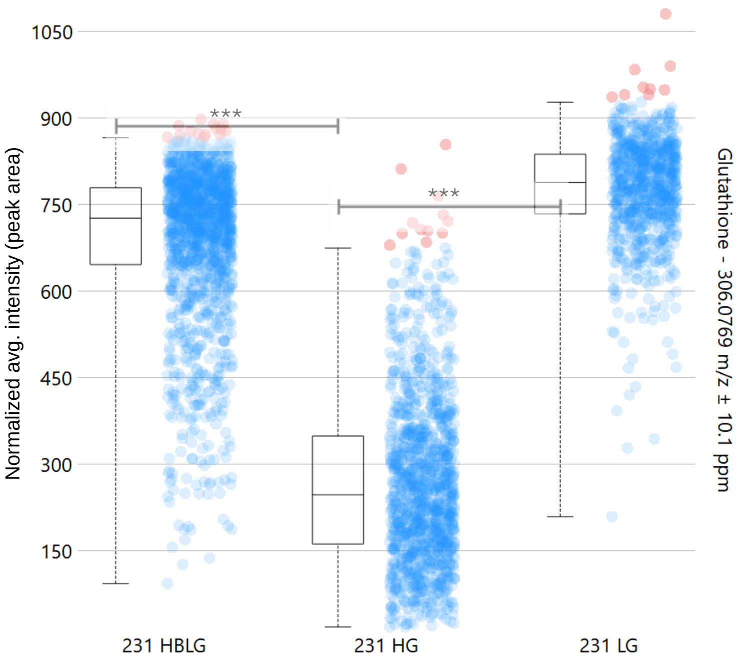


**Supplementary Figure S21.** Relative distributions of Glutathione in MDA-MB-231 cells grown in different nutrient media, HG, LG and HBLG. Root mean square normalization was followed for computing average intensities and Wilcoxon rank sum corrected p values were used for assessing statistical significance where *** indicates $p < 0.001$.

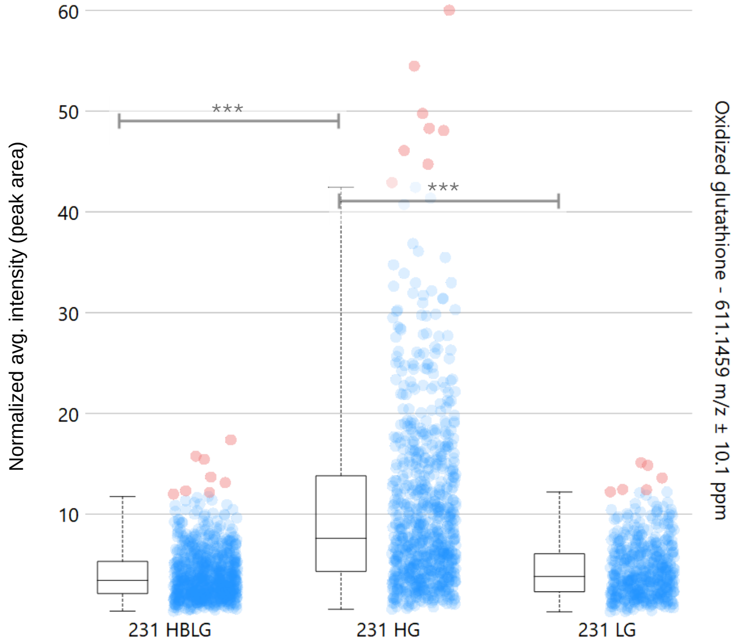


**Supplementary Figure S22.** Relative distributions of Oxidized glutathione in MDA-MB-231 cells grown in different nutrient media, HG, LG and HBLG. Root mean square normalization was followed for computing average intensities and Wilcoxon rank sum corrected p values were used for assessing statistical significance where *** indicates $p < 0.001$.

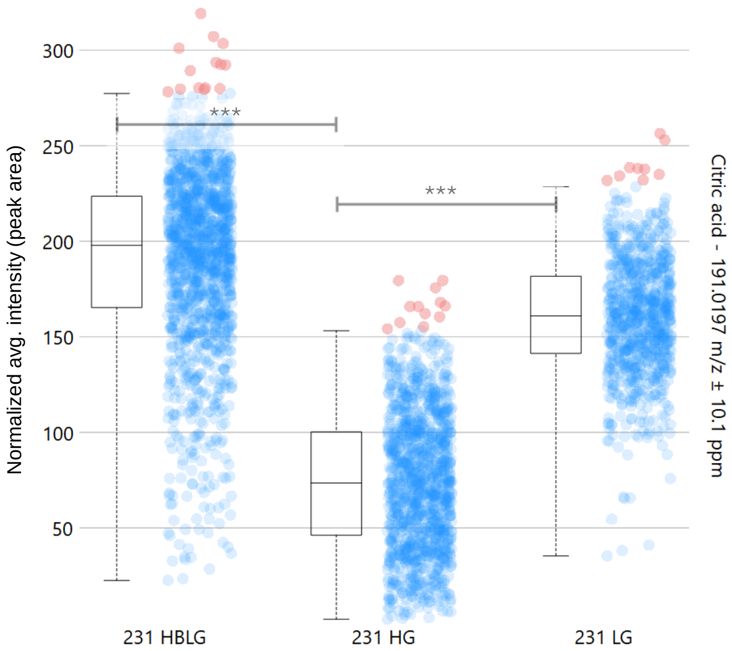


**Supplementary Figure S23.** Relative distributions of Citric acid in MDA-MB-231 cells grown in different nutrient media, HG, LG and HBLG. Root mean square normalization was followed for computing average intensities and Wilcoxon rank sum corrected p values were used for assessing statistical significance where *** indicates $p < 0.001$.

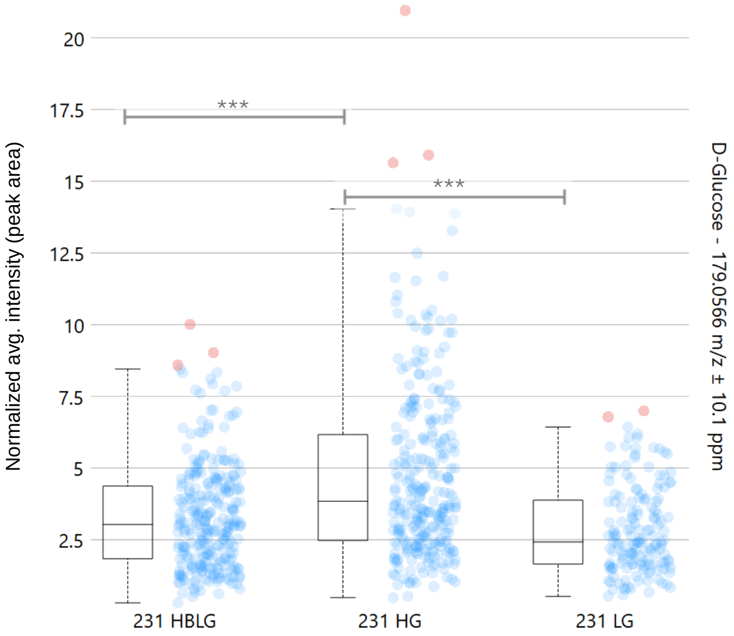


**Supplementary Figure S24.** Relative distributions of D-glucose in MDA-MB-231 cells grown in different nutrient media, HG, LG and HBLG. Root mean square normalization was followed for computing average intensities and Wilcoxon rank sum corrected p values were used for assessing statistical significance where *** indicates $p < 0.001$.

**Supplementary Table S1.** List of primers used for rt-*q*PCR studies

| Name | Sequence of primer (5'-3') |
|---|---|
| h-ST6GAL1-F | GCAAAGATCAGAGTGAAACAG |
| h-ST6GAL1-R | CACCTCATCGCAGACATG |
| h-GFPT1-F | TTGCCTGTGATGGTGGAACT |
| h-GFPT1-R | GTGATATGGAACTGCCAACTGT |
| h-GNE-F | GAAGCATACGCCTCTGGAATGG |
| h-GNE-R | CAGCCTCATCTTTTGGCACTGAC |
| h-FASN-F | CATCCAGATAGGCCTCATAGAC |
| h-FASN-R | CTCCATGAAGTAGGAGTGGAAG |
| h-ACTB-F | TGACGTGGACATCCGCAAAG |
| h-ACTB-R | CTGGAAGGTGGACAGCGAGG |
